# High throughput imaging identifies a spatially localized response of primary fetal pulmonary artery endothelial cells to insulin-like growth factor 1 treatment

**DOI:** 10.1101/674499

**Authors:** Christina Kim, Gregory J Seedorf, Steven H Abman, Douglas P Shepherd

**Affiliations:** Department of Surgery, University of Colorado Anschutz Medical Campus, Aurora, CO 80045; Pediatric Heart Lung Center, University of Colorado Anschutz Medical Campus, Aurora, CO 80045; Department of Pediatrics, University of Colorado Anschutz Medical Campus, Aurora, CO 80045; Department of Pharmacology, University of Colorado Anschutz Medical Campus, Aurora, CO 80045

**Keywords:** High throughput imaging, spatial analysis, drug response

## Abstract

A common strategy to measure the efficacy of drug treatment is the *in vitro* comparison of ensemble readouts with and without treatment, such as proliferation and cell death. A fundamental assumption underlying this approach is there is minimal cell to cell variability in the response to drug. Here, we demonstrate that ensemble and non-spatial single cell readouts applied to primary cells lead to incomplete conclusions due to cell to cell variability. We exposed primary fetal pulmonary artery endothelial cells (PAEC) isolated from healthy newborn healthy and persistent pulmonary hypertension of the newborn (PPHN) sheep to the growth hormone insulin-like growth factor 1 (IGF-1). We found that IGF-1 increased proliferation and branch points in tube formation assays but not angiogenic signaling proteins at the population level for both cell types. We hypothesized that this molecular ambiguity was due to the presence of cellular subpopulations with variable responses to IGF-1. Using high throughput single cell imaging, we discovered a spatially localized response to IGF-1. This suggests localized signaling or heritable cell response to external stimuli may ultimately be responsible for our observations. Discovering and further exploring these rare cells is critical to finding new molecular targets to restore cellular function.

## 1. Introduction

Drug discovery often relies upon initial results from treating isogenic cell lines that mimic the phenotype for the disease of interest.^1–4^ However, there is growing evidence that intrinsic and extrinsic fluctuations can generate numerous unique cell states within isogenic cell lines.^5–7^ There is also evidence that after controlling for cell cycle and cell state, the remaining fluctuations are solely due to biochemical reaction noise.^8,9^ Recent studies have demonstrated that variability is conferred from mother to siblings cells,^5^ inheritable fluctuations in gene expression found within clonal cancer cell lines may lead to drug resistance,^6^ and distributions in the number of mitochondria can explain cell death due to TNF-related apoptosis.^7^

Regardless of the underlying mechanism, cell to cell variability has broad implications outside of studying mechanisms heterogeneity in clonal cell lines and drug resistance in cancer. For example, in the case of developmental diseases of the newborn, changes to the external environment during gestation leads to altered cellular genotype and phenotype that ultimately lead to a sustained disease phenotype. Clinically, there is a stagnation of effective therapies for many developmental diseases.^10^ One contributing factor is the growing number of promising *in vitro* studies of drug therapies that fail to translate to meaningful results in clinical trials.^11^ While there are many possible explanations for the failure of clinical trials, one potential explanation is that immortalized cell lines fail to represent the disease phenotype.^2,12^ This failure has led to a new emphasis on performing drug discovering using primary cells isolated from verified disease models.^1^ However, in non-clonal primary cells, single-cell heterogeneity could be due to multiple epi-genetic populations, variability due to a stochastic response to fast environmental response changes, or other forms of variability that are just beginning to be explored.^13–15^ This additional heterogeneity necessitates a careful experimental approach that integrates traditional ensemble readouts with high-throughput single-cell measurements of molecular signaling (Figure 1).

**Figure 1.**
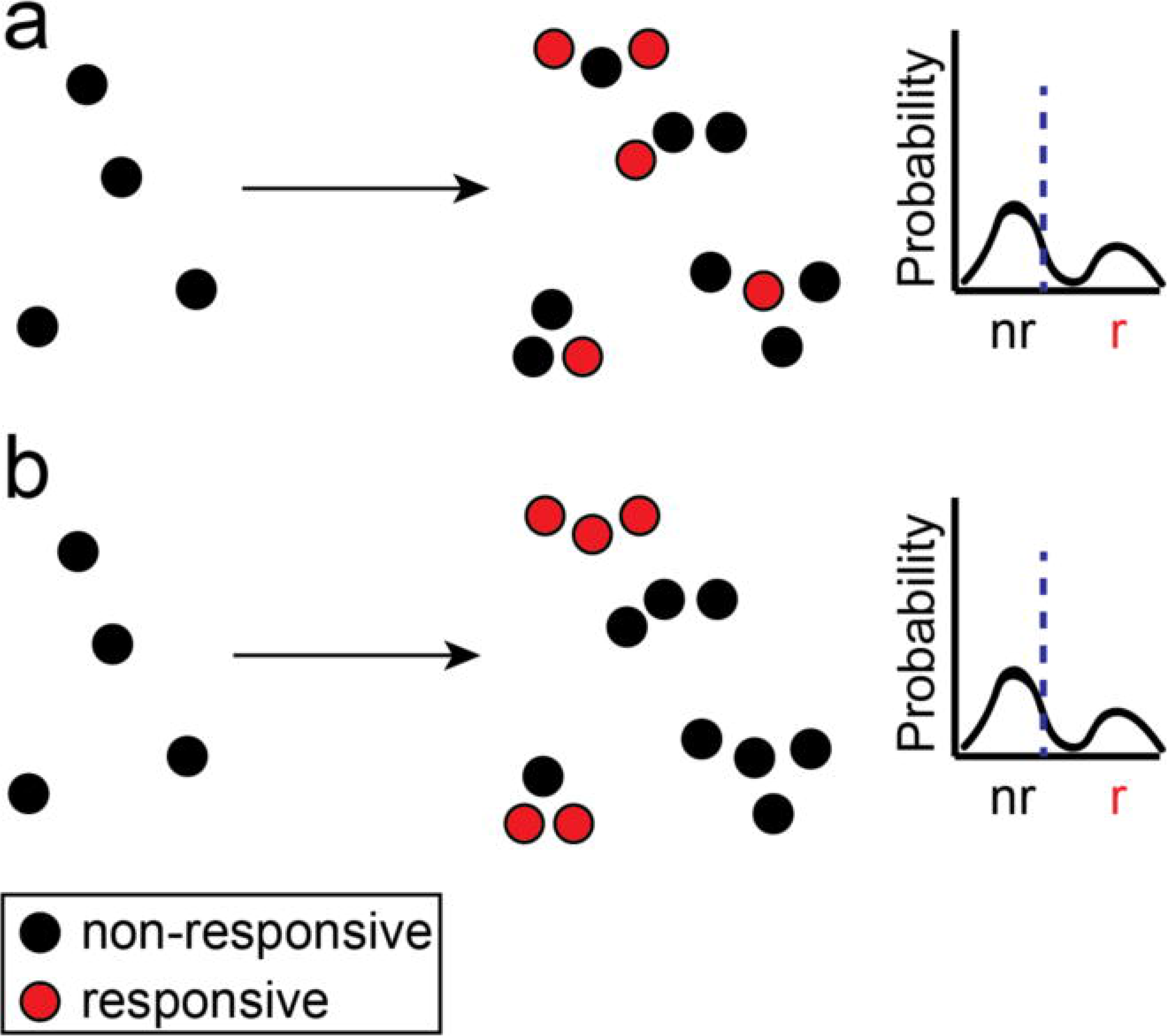
Spatially associated drug responsiveness. Single-cell measurements of drug responsiveness (a) without spatial information or (b) with spatial information will provide the same mean (blue dashed line) and single-cell distributions. Only spatially resolved measurements can determine if the spatial distribution of responsive cells is random or correlated.

One developmental disease with a clinical stagnation of new molecular therapies is persistent pulmonary hypertension of the newborn (PPHN). PPHN represents the failure of the lung circulation to adapt to postnatal conditions in term and preterm infants and can be associated with several cardiopulmonary diseases. PPHN is manifested by elevated pulmonary vascular resistance (PVR) after birth that leads to profound hypoxemia due to right-to-left extrapulmonary shunting. High PVR can be due to elevated pulmonary vascular tone, hypertensive remodeling of the vessel wall and decreased vascular growth due to impaired angiogenesis. Diverse mechanisms potentially impair vasodilation at birth, but past experimental and clinical studies have shown that decreased production of nitric oxide (NO) contributes to high PVR at birth, and that inhaled NO has been FDA-approved for the management of PPHN in the setting of acute respiratory failure. Although inhaled NO improves oxygenation, lowers PVR and reduces the need for ECMO and death in PPHN, some infants fail to respond to therapy, suggesting the need for additional therapies.^16^

A potential new molecular target for PPHN therapy is the insulin-like growth factor-1 (IGF-1) pathway. IGF-1 is a potent growth hormone that contributes to normal lung development and is critical for endothelial growth and survival.^17^ Previous studies have shown that a relative deficiency in IGF-1 in preterm infants can affect multiple organ systems including the brain, eyes, lungs and cardiovascular system, contributing to diseases including retinopathy of prematurity (ROP) and BPD.^18–20^ Paradoxically, an increase in IGF-1 expression has been associated with pulmonary hypertension (PH) due to chronic hypoxia in rodents and calves, which was considered to be caused by smooth muscle cell proliferation.^17,21^ The effects of IGF-1 in the setting of PPHN remains controversial and whether IGF-1 can restore endothelial cell function is unknown.

In this study, we aim to test if IGF-1 administration can improve the function of diseased endothelial cells. Instead of isogenic and immortalized endothelial cells, we utilized primary fetal pulmonary artery endothelial cells (PAEC) isolated from normal fetal sheep (normal PAEC) and fetal sheep with severe intrauterine PH (PPHN PAEC) (Figure 2). Past work has shown that ductus arteriosus ligation in utero in late gestation lambs provides a useful model for studies of PPHN.^22^ In this model, intrauterine PH due to ductus ligation impairs vascular growth, increases pulmonary vascular resistance and causes sustained PPHN at birth.^23,24^ Previous *in vitro* studies have shown that primary PPHN PAECs had impaired growth and tube formation and decreased expression of pro-angiogenic genes.^25^

**Figure 2.**
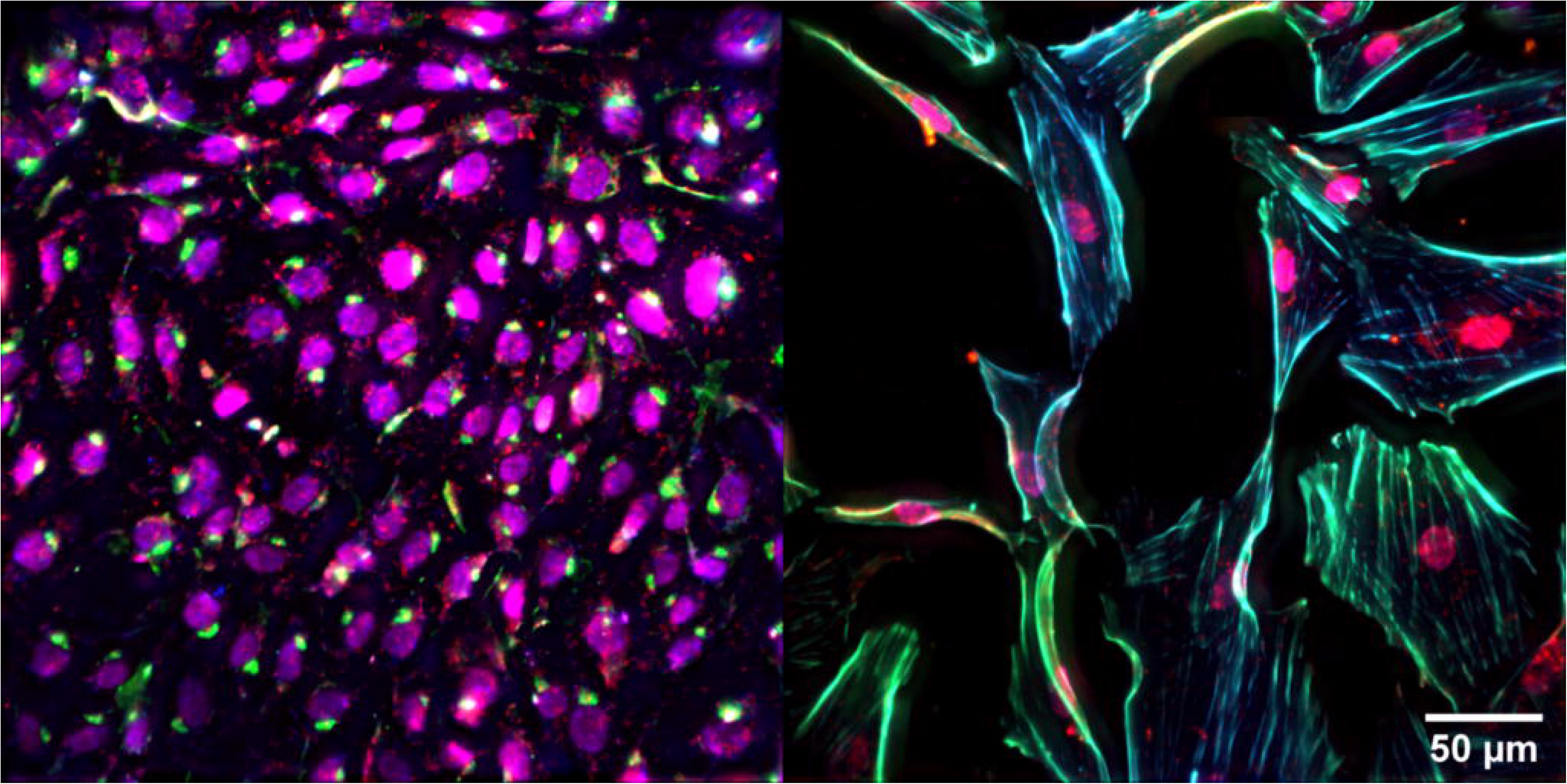
Primary pulmonary artery endothelial cells. Representative images of nuclei (magenta), actin (blue), eNOS (green), VEGF (red) in (left) normal PAEC and (right) PPHN PAEC.

Based on the results that IGF-1 deficiency can alter lung and vascular development, we hypothesized that IGF-1 treatment would increase PPHN PAEC proliferation, tube formation, and increase the production of two downstream proteins associated with angiogenesis, vascular endothelial growth factor (VEGF) and endothelial nitric oxide synthase (eNOS). We tested this hypothesis utilizing standard ensemble assays and found that IGF-1 increases proliferation and branch point in tube formation assay for normal and PPHN PAEC. However, we found no evidence for changes in VEGF and eNOS at the ensemble level. These findings led us to hypothesize that there exist distinct sub-population of PAEC with differing molecular responses to IGF-1 administration. Using high-throughput single-cell fluorescence imaging, we quantified the actin cytoskeleton, VEGF and eNOS expression, and new protein synthesis in over 1 million individual PAEC in snapshots across cell type, drug administration, and time points. This high-density imaging dataset confirmed the existence of multiple PAEC sub-populations. Further analysis of these data provided evidence that PAEC sub-populations are spatially correlated, suggesting a localized response to IGF-1 administration.

Our findings suggest that there exist subtle variations across multiple subpopulations within PAEC, hidden by common ensemble measurement methods. One or more of these variations lead to a spatially associated response to IGF-1 administration. This spatial association is sustained during IGF-1 administration in PPHN PAEC as compared to normal PAEC. More broadly, our findings suggest that exploiting the spatial correlations in single-cell imaging data sets will provide new insight into how molecular therapies interact with target cells.

## 2. Methods

### 2.1 Isolation and cell culture of fetal ovine PAECs

All procedures and protocols were reviewed and approved by the Animal Care and Use Committee at the University of Colorado Anschutz Medical Campus. The left and right pulmonary arteries were isolated from late-gestation normal fetal sheep (mixed-breed Columbia-Rambouillet pregnant ewes at 135 days gestation) and from fetal sheep that had undergone partial ligation of the ductus arteriosus in utero 7-10 days before euthanasia, as previously described).^25^ Proximal PAECs were isolated as previously described.^25^ Briefly, proximal pulmonary arteries were separated from fetal sheep, and branching vessels were ligated. Collagenase was used to separate endothelial cells from the vessel wall. PAECs were plated and grown in Dulbecco’s modified Eagle medium (DMEM) and 10% fetal bovine serum (FBS) (10-013 corning cellgro, Manassas VA, Gemini Bio-Products, Sacramento CA). Endothelial cell phenotype was confirmed by the typical cobblestone appearance and von Willebrand Factor expression (A0082 Dako-Agilent Santa Clara CA) (Supplemental Figure 1). Cells from passages 4 through 7 were used for the experiments and cells from each animal were kept separate throughout all passages and experiments.

### 2.2 IGF-1 compound

Shire Pharmaceutics provided Insulin-like growth factor 1 with binding protein 3 (IGF-1). A dose-response with IGF-1 was used to determine the most responsive dose of IGF-1 with normal PAECs (Supplemental Figure 2).

### 2.3 Proliferation assays

The effects of IGF-1 on PAEC growth was compared between normal and PPHN PAECs. 60,000 cells were plated per well and adhered overnight. Media was then changed to IGF-1 at 250 ng/ml in 2.5% FBS in DMEM (treatment) or DMEM alone (vehicle). Media was changed daily, and cell counts were performed daily, up to day 3.

### 2.4 Tube formation assays

The ability of fetal PAECs to migrate and form multicellular structures *in vitro* was assessed by plating PAECs on collagen. PAECs were placed on collagen plated wells at a density of 50,000 cells/well in DMEM supplemented with 0.5% FBS and DMEM supplemented with 0.5% FBS and 250 ng/ml of IGF-1. PAECs were incubated for 18-24 hours for maximal tube formation, fixed with 4% paraformaldehyde (PFA) (47608, MilliporeSigma, St. Louis, MO) and branch points were counted from two different locations per well.

### 2.5 Western Blot assays

PAEC from normal and PPHN lambs were grown on 150-mm cloning plates with DMEM and 10% FBS. When the cells reached 80-90% confluence, the cell lysates were collected. The protein content of samples was determined using the BCA protein assay (23225, Pierce Biotechnology, Rockford, IL), using bovine serum albumin as the standard. A 25 μg protein sample was added to each lane and resolved by sodium dodecyl sulfate-polyacrylamide gel electrophoresis. Proteins from the gel were then transferred to a nitrocellulose membrane. VEGF (sc-152, Santa Cruz Biotechnology, Santa Cruz, CA), eNOS (610297, BD Biosciences, San Jose, CA) and β-actin (A2228, Sigma, St. Louis, MO) were detected using appropriate normals and molecular weight as identified by the manufacturer for each protein of interest.

### 2.6 Proliferation, tube formation, and Western Blot assay analysis

Proliferation, branch point, and western blot analyses were analyzed using Python 3.6. The non-parametric Kruskal-Wallis test was applied to determine statistical significance for proliferation, tube formation assays and western blot analysis.

### 2.7 Immunofluorescence assays

PAEC from normal and PPHN lambs were grown on gelatin plated glass slides (No. 1.5, 22×22 mm, Arthur H. Thomas Company, Philadelphia, PA) and grown to 70-85% confluence. These PAECs were then serum deprived in untreated DMEM for 24 hours. The PAECs were then subsequently treated with 2.5% FBS in DMEM or 2.5% FBS in DMEM with 250 ng/ml of IGF-1 for specific time points of 0, 15, 30 minutes, 1, 2, 4 and 24 hours. The slides were then fixed with 4% PFA (47608, MilliporeSigma, St. Louis, MO) for 15 minutes, washed with PBS 3 times and then permeabilized with Triton X-100 at 0.25% in PBS for 10 minutes. The PAECs were then blocked with 1% BSA in PBST for 30 minutes and incubated in primary antibodies: VEGF (1:250, sc-152, Santa Cruz Biotechnology, Dallas, TX) and eNOS (1:250, 610297, Santa Cruz Biotechnology, Dallas, TX), overnight. To prevent bleaching, the subsequent steps were performed under low light. The PAECs were incubated in secondary antibody: Alexa Fluor 555 (A-31570, ThermoFisher Scientific, Waltham, MA), Alexa Fluor 647 (A-31574, ThermoFisher Scientific, Waltham, MA) and subsequently stained with acti-stain 488, Phalloidin (PHDG1, Cytoskeleton, Inc., Denver, CO) and NucBlue Fixed Cell Stain ReadyProbes reagent (R37606, Life Technologies, Carlsbad, CA). The glass slides were plated on standard microscope slides with 15-20 μl of GLOX buffer + enzyme (1 μl of 3.7 mg/ml glucose oxidase (G7141, MilliporeSigma, St. Louis, MO), 1 μl catalase (C30, MilliporeSigma, St. Louis, MO)) and sealed with nail polish. In addition, separate experiments with von Willebrand factor (1:250, sc-8068, Santa Cruz Biotechnology, Santa Cruz, CA) were performed with Alexa Fluor 647 (A-21447, ThermoFisher Scientific, Waltham, MA), acti-stain 488, Phalloidin (PHDG1, Cytoskeleton, Inc., Denver, CO) and NucBlue Fixed Cell Stain ReadyProbes reagent (R37606, Life Technologies, Carlsbad, CA) and fixed as above.

### 2.8 Protein translation assays

PAECs were grown to 70-80% confluence on glass coverslips (#1.5, 22 mm × 22 mm) and treated with 2.5% FBS or 2.5% FBS and 250 ng/ml of IGF-1. Samples were then treated with Click-iT HPG (Component A) (Click-iT HPG Alexa Fluor Protein Synthesis Assay, C10428, C10429, Life Technologies, Carlsbad, CA), an amino acid analog of methionine that integrates into newly formed protein, per the drug pre-incubation protocol and fixed at 0, 1, 24 hours. Click-iT reaction cocktail was prepared per protocol and slides were incubated as outlined, DAPI was added to identify individual cells and slides were placed on microscope slides and fixed as above.

### 2.9 Fluorescence microscopy

All imaging was performed utilizing a homemade structured illumination microscope built on an Olympus IX71 microscope body (Olympus Corporation, Center Valley, PA). In this work, we did not utilize the structured illumination capabilities of our instrument because diffraction limited imaging is sufficient for nuclear and cytosolic protein quantification. Therefore, the instrument was run in epi-fluorescence mode followed by deconvolution.

An LED light source (Spectra-X, Lumencor, Beaverton, OR) equipped with a multi-emitter specific filter set (LED-DA/FI/TR/Cy5-4X-A, Semrock, Lake Forest, IL) was coupled to a 3 mm liquid light guide (LLG). The LLG was coupled into a multi-element collimator (LLG3A6, Thorlabs, Newton, NJ). Collimated light was directed onto a digital micromirror device (DMD, DLP6500, Texas Instruments, Dallas, TX) at a 22-degree angle to the horizontal. In our configuration, DMD *off* pixels are directed on axis into the excitation light path and DMD *on* pixels are directed off axis. The pattern displayed on the DMD was relayed to the sample plane using a 2-inch achromatic doublet (AC508-300-A-ML, Thorlabs, Newton, NJ) and an oil immersion objective (UPLFLN 40XO, Olympus Corporation, Center Valley, PA) mounted on an objective piezo (FN200, Mad City Labs, Madison, WI). This gives an effective DMD pixel size of 112 nm. To ensure correct alignment of the excitation arm, the DMD was positioned using a translation stage (PT1, Thorlabs, Newton, NJ), a rotation stage (PR01, Thorlabs, Newton, NJ), and tilt stage (FP90, Thorlabs, Newton, NJ) using a 3D printed optic mount. The DMD was then positioned along and around the microscope optical axis to achieve the highest modulation of a checkerboard pattern, built into MicroManager 2.0 gamma (MM 2.0), on a homemade Fluorescein slide. ^26,27^ A one to one mapping of DMD pixels to camera pixels was carried out using SIMToolbox. ^28,29^ Samples were mounted on an XY translation stage (Microdrive, Mad City Labs, Madison, WI). Fluorescence was collected through the oil immersion objective, passed through the dichroic and quad-band emission filter (LED-DA/FI/TR/Cy5-4X-A, Semrock, Lake Forest, IL), and exited the microscope body. The tube lens and all windows were removed from the microscope body. Fluorescence passed through the microscope tube lens (SWTLU-C, Olympus Corporation, Center Valley, PA), a dichroic beam splitter (FF562-Di03-25×36, Semrock, Lake Forest, IL) housed in a kinematic dichroic mount (DFM1, Thorlabs, Newton, NJ) and was directed onto two sCMOS cameras (OrcaFlash4.0 v2, Hamamatsu Corporation, Bridgewater, NJ). The short wavelength camera was mounted on an adjustable rotation mount (LCP02R, Thorlabs, Newton, NJ). The long wavelength camera was mounted on an adjustable XYZ mount (CXYZ1, Thorlabs, Newton, NJ). This enabled simultaneous and aligned dual-color imaging. The effective pixel size at each camera is 162.5 nm, roughly ⅔ larger than the Nyquist limit for the detection objective. This trade-off was necessary to maintain a large FOV (332.8 × 332.8 um). Chromatic alignment, spectral cross-talk quantification, and defocus compensation of the two cameras was performed by the user at the pixel level using 100 nm multicolor beads (T14792, Life Technologies, Carlsbad, CA).

All electronics were connected using USB3/3.1, except for HDMI for pixel display on the DMD device, to a Windows 7 PC with 32 GB of RAM and 2 Tb of SSD storage. MM 2.0 was used to set up and acquire all acquisitions.^26,27^ Epifluorescence data was collected with all DMD pixels set to *off* (corresponding to light being directed into the excitation optics) using the projector plugin, autofocus plugin, and multi-dimensional acquisition normal in MM 2.0. 100 image areas were acquired for each slide in a 10×10 grid with no overlap. For each image area within the grid, three or four axial stacks with independent excitation/emission wavelengths were acquired. Individual image stacks consisted of 58 axial steps, with an axial step size of 350 nm, and an XY pixel size of 162.5 nm. Raw 16-bit images were transferred to a Ubuntu 18.04 LTS server with 24 cores, 128 GB of RAM, GPUs (2x TITAN-X, Nvidia, Santa Clara, CA) and 60 TB of high-speed storage for processing. All raw 16-bit images were corrected for CMOS specific camera noise,^30^ deconvolved on a GPU using measured point spread functions (PSF) (Microvolution, Palo Alto, CA), and flat-field corrected.^31,32^

### 2.10 Image quantification

Deconvolved and flat-field corrected image stacks were split into individual channels and maximum projected using Fiji.^32^ CellProfiler 3.1 was used to segment nuclei boundaries, construct cell boundaries, filter out identified cells without nuclei, and filter out identified cells not meeting a user-set size threshold.^33^ For each imaging set, the nuclear segmentation and filtering options were manually verified and refined as needed before batch quantification. The same CellProfiler 3.1 pipeline was used to quantify molecular label intensity, cell morphology, texture, and adjacency for all identified and accepted cells. 389 measurements were exported for each identified nucleus and cell. Single cell measurements were exported as a text file for further analysis.

### 2.11 Single-cell analysis

Data were imported into Python 3.6 as a Pandas data structure.^34^ Marginal single-cell distributions were tested for significance using the non-parametric Kruskal-Wallis test. Spatial analysis of normal and PPHN PAEC experiments were conducted separately due to the observed differences in cell morphology. The median absolute deviation (MAD) was calculated for every feature in normal PAEC at t=0 and PPHN PAEC at t=0. Those features with MAD=0 were removed from the feature set. For all experiments, each feature was then normalized by subtracting the median and dividing by 1.4826*MAD.^35^ Principal component analysis (PCA) was then used to determine the number of principal components required to capture 99% of the variance for all normal PAEC conditions and all PPHN PAEC conditions. For both cell types, this required 119 principal components. We then calculated the null distribution of single-cell correlations for all conditions by repeatedly randomly selecting 2 cells from each condition and calculating the Pearson correlation coefficient. We repeated this calculation for 2000 random draws and then calculated the median Pearson correlation coefficient. We then calculated spatial single-cell correlations by calculating the median Pearson correlation for the n=20 nearest neighbors for all cells.

### 2.12 Data availability

Code to generate all analyses in the paper are provided as supplementary material. Data used to generate the figures in the main text and supplement can be found at ref. ^49^

## 3. Results and Discussion

### 3.1 Population response to IGF-1 administration

We first asked if IGF-1 administration could increase three common experimental readouts of improved endothelial cell function: proliferation, branch points in a tube formation assay, and angiogenic signaling. Based on a dose-response study, we determined the maximum increase in proliferation occurred at 250 ng/ml of IGF-1 in both normal and PPHN PAEC (Supplemental Figure 2).

IGF-1 administration over three days increased the growth of normal and PPHN PAEC growth by 32% and 51%, respectively (Figure 3a). We performed additional normals by quantifying PAEC proliferation in response to IGF-1 treatment after inhibiting the IGF-1 receptor (IGF-1R) and mTORC1 (Supplemental Figures 3 and 4). We found that inhibiting IGF-1R blocked the observed PAEC response to IGF-1 while inhibiting mTORC1 does not block observed PAEC response to IGF-1.

**Figure 3.**
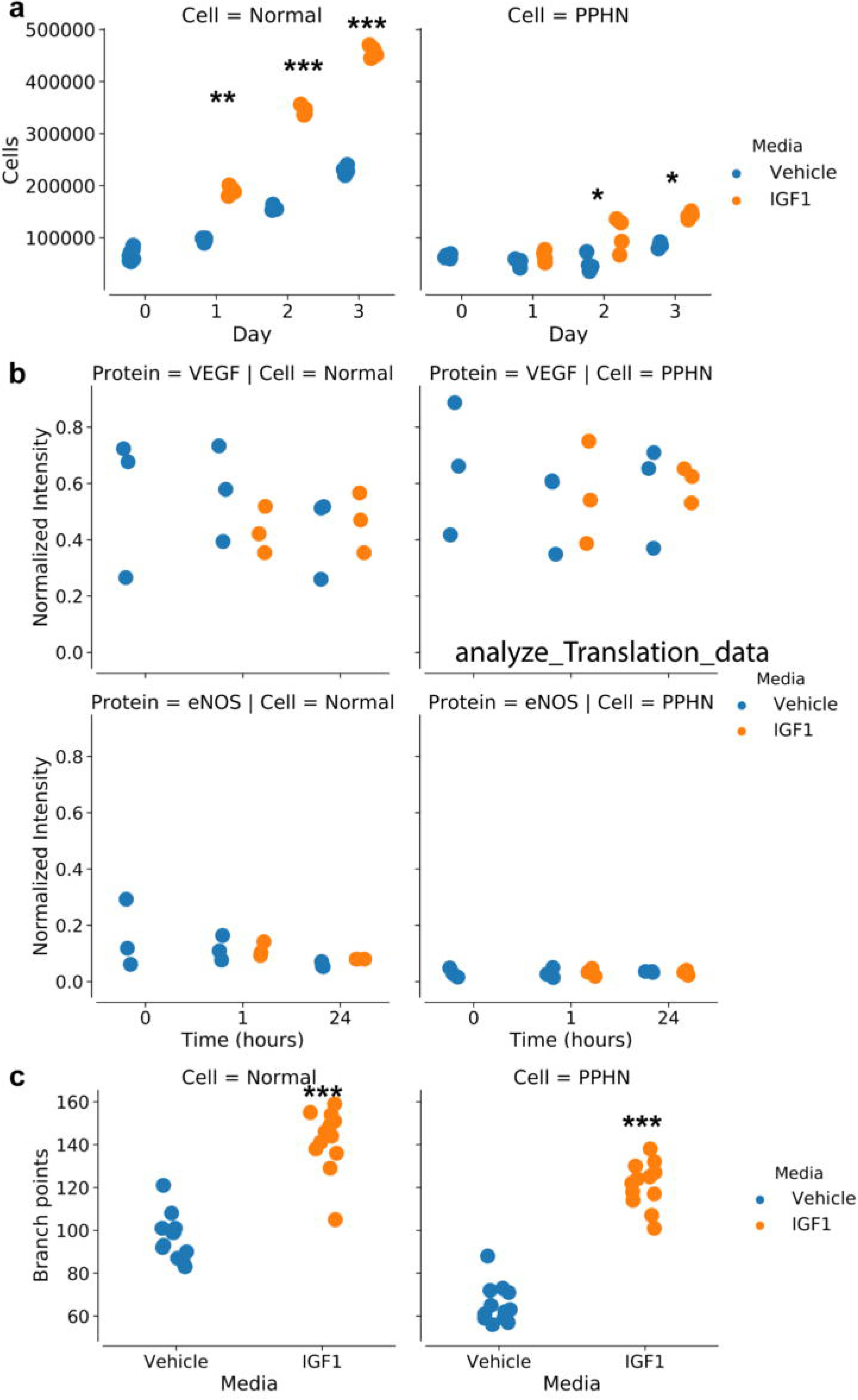
Population response to IGF-1 administration. a) Proliferation for normal PAEC and PPHN PAEC with vehicle (blue) or with IGF-1 administration (orange) over three days. b) Western blot analysis for VEGF protein expression (n=3, normalized to β-actin). d) Western blot analysis for eNOS protein expression (n=3, normalized to β-actin). c) Branch points increase for normal PAEC and for PPHN PAEC upon IGF1 administration. All comparisons performed using the Kruskal-Wallis test. * p<.01; ** p<.001; **** p<.0001. Non-significant test results (greater than p>.05) are not demarcated.

**Figure 4.**
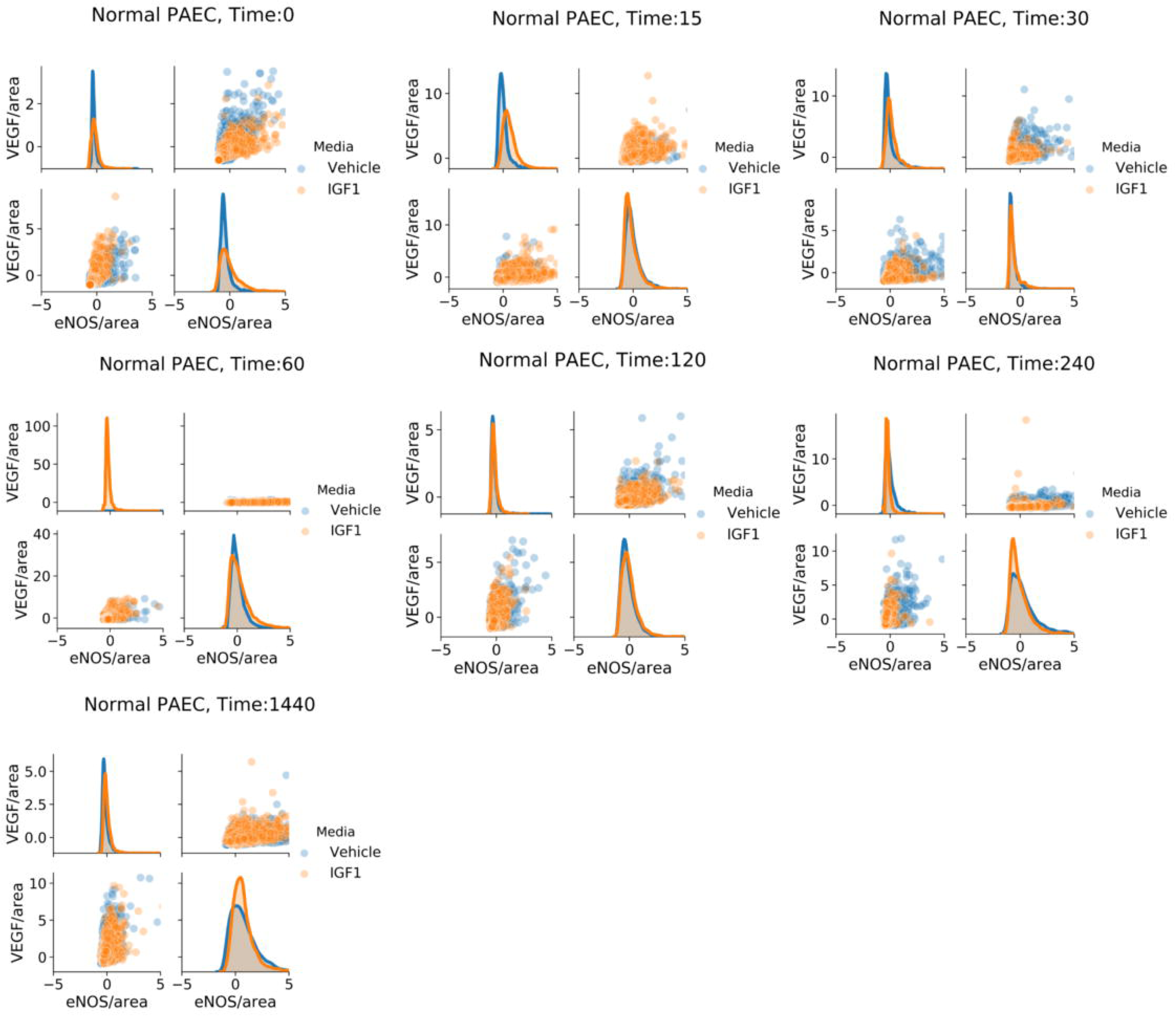
Normal PAEC marginal and joint distributions of VEGF and eNOS production. For all cell and time snapshots, the normalized marginal probability distributions were plotted for both vehicle (blue) and IGF-1 (orange) media. All marginal distribution tests performed using the Kruskal-Wallis test. * p<.01; ** p<.001; **** p<.0001. Non-significant test results (greater than p>.05) are not demarcated.

To quantify changes in angiogenic signaling, we performed western blot analysis for VEGF and eNOS protein content from normal and PPHN PAEC lysates at multiple time points following treatment with vehicle (normal) or IGF-1 (t = 0, 1, 2, 4, 24 hours). We found decreased eNOS protein content in PPHN PAEC in comparison with normals under basal conditions. However, IGF-1 treatment did not affect VEGF or eNOS protein expression in normal or PPHN PAECs within each group (Figure 3b, Supplemental Figure 5).

**Figure 5.**
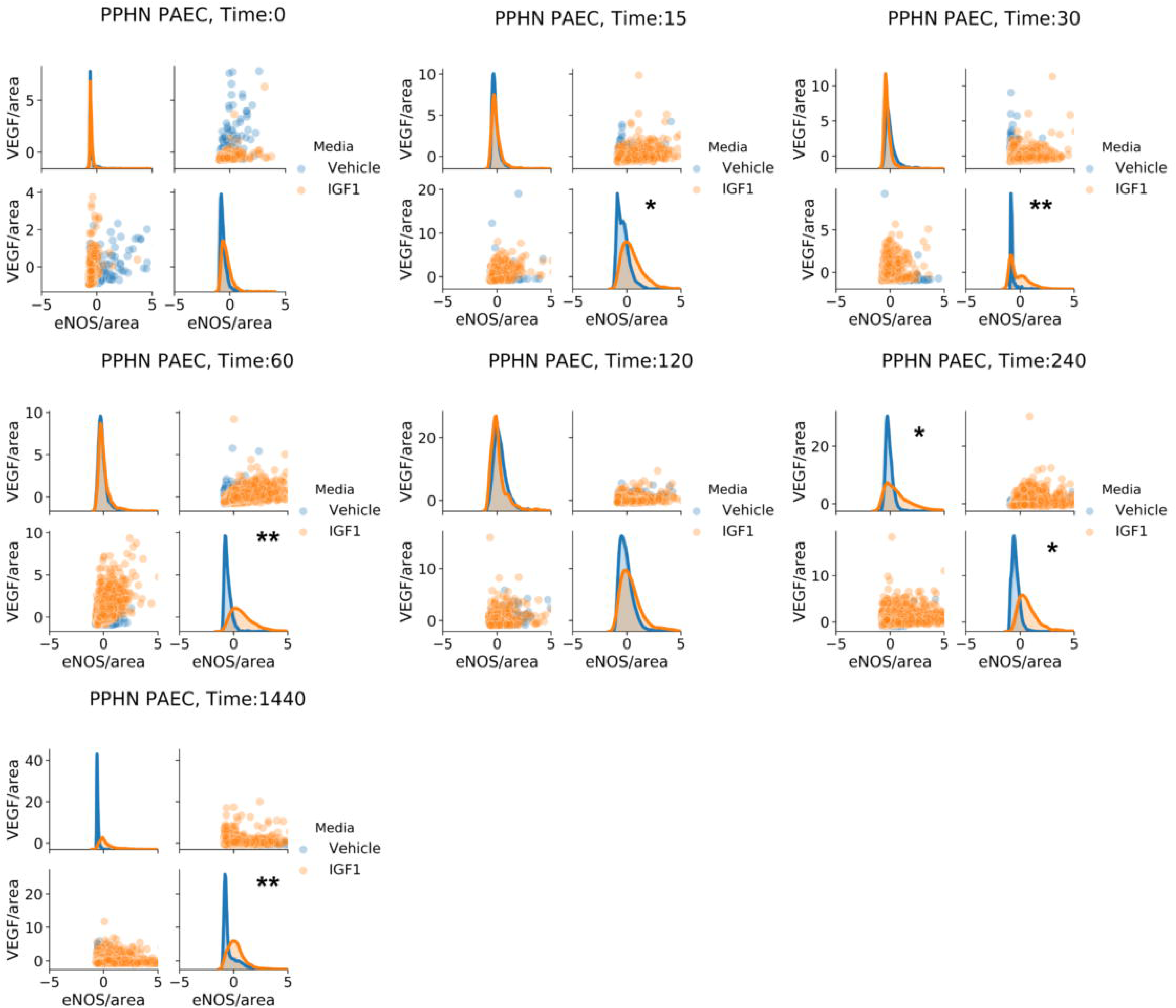
a) PPHN PAEC marginal and joint distributions of VEGF and eNOS production. For all cell and time snapshots, the normalized marginal probability distributions were plotted for both vehicle (blue) and IGF-1 (orange) media. All marginal distribution tests performed using the Kruskal-Wallis test. * p<.01; ** p<.001; **** p<.0001. Non-significant test results (greater than p>.05) are not demarcated.

Tube formation assays evaluate the ability of endothelial cells *in vitro* to migrate and form multicellular structures, which is a surrogate for endothelial cells function.^36^ Normal PAECs had a 47% increase with IGF-1 treatment. PPHN PAECs in vehicle media had a 47% decrease in branch points compared to normal PAECs. With IGF-1 treatment there was an 85% increase in branch points in PPHN PAECs (Figure 3c).

These results demonstrated that IGF-1 administration restored some functionality of PPHN PAEC despite the drastically different initial cellular phenotype as compared to normal PAEC. The increase in proliferation and branch points was less pronounced for PPHN PAEC than normal PAEC. Typically, these results would lead to the conclusion that IGF-1 was partially effective at globally restoring endothelial cell function. Our single-cell studies revealed this is an incorrect conclusion of the cellular response to IGF-1 administration.

### 3.2 Single-cell response to IGF-1 administration

We next asked if the difference in response between normal and PPHN PAEC to IGF-1 administration may be due to multiple PAEC sub-populations. To answer this question, we utilized high-throughput single-cell immunofluorescence imaging to quantify VEGF, eNOS, and total new protein synthesis in over 1 million individual PAEC across cell type, treatment type, and time post-administration. This density ensured that we sufficiently sampled the positive, asymmetric distribution of single-cell response to IGF-1 so as to not to bias our results.^15,37^

We found that IGF-1 administration changed the shape of area normalized single-cell distributions of VEGF and eNOS for both normal and PPHN PAEC (Figures 4 and 5). However, we did not find evidence of distinct subpopulations and instead observed a continuum of expression. We additionally found that IGF-1 administration did not change the shape of area normalized single-cell distributions of total protein expression for either normal or PPHN PAEC (Figure 6).

**Figure 6.**
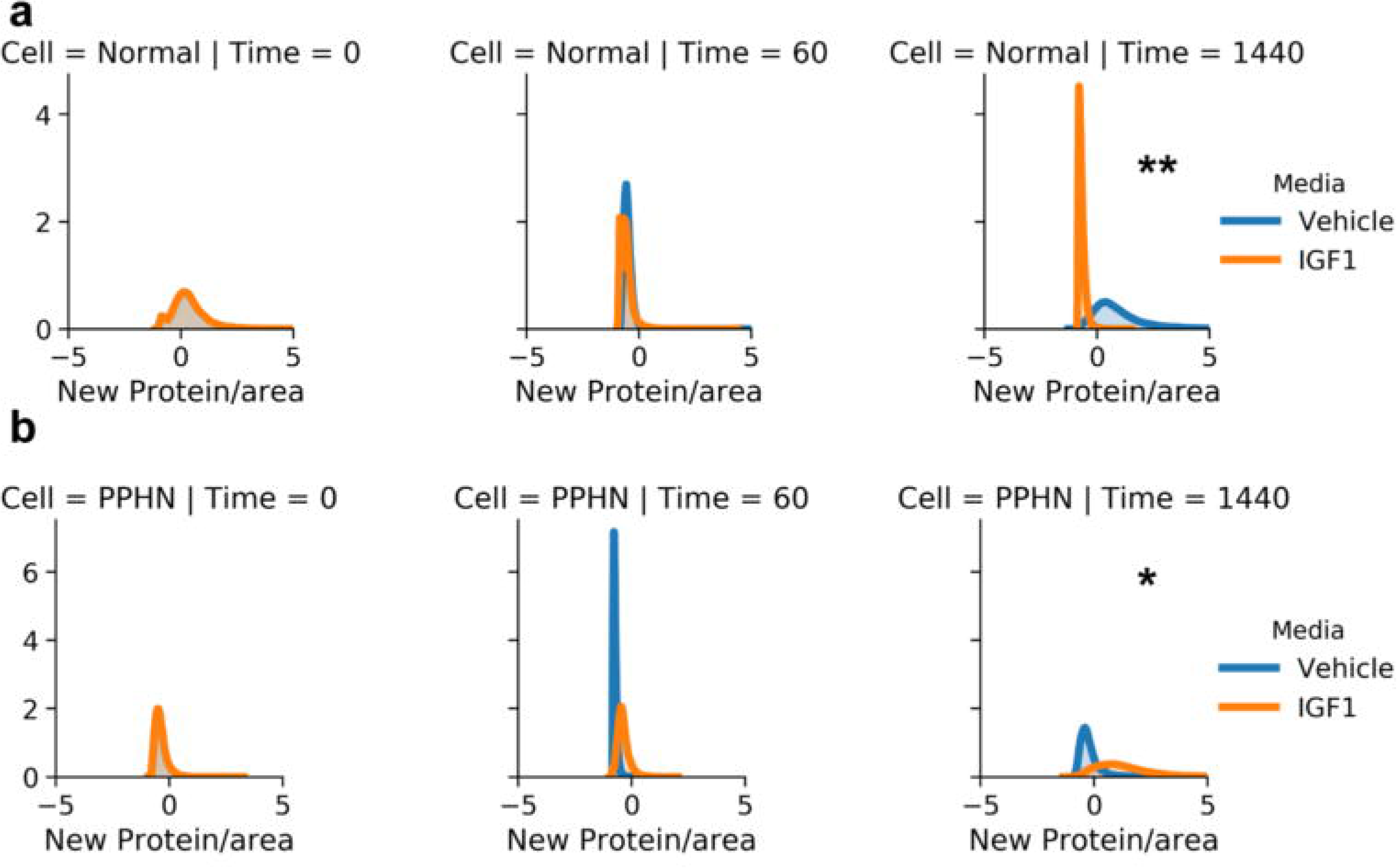
a) Normal PAEC marginal and joint distributions of total protein production. For all cell and time snapshots, the normalized marginal probability distributions were plotted for both vehicle (blue) and IGF-1 (orange) media. b) PPHN PAEC marginal and joint distributions of total protein production. For all cell and time snapshots, the normalized marginal probability distributions were plotted for both vehicle (blue) and IGF-1 (orange) media. All marginal distribution tests performed using the Kruskal-Wallis test. * p<.01; ** p<.001; **** p<.0001. Non-significant test results (greater than p>.05) are not demarcated.

These results demonstrated that IGF-1 treatment induced a diverse response in normal and PPHN PAEC. One potential explanation is that specific PAEC responded to IGF-1 and those PAEC were responsible for the observed increase in proliferation. However, our single-cell results were not conclusive based on the limited quantification of VEGF and eNOS expression. Beyond VEGF and eNOS expression, our data contained information on cell morphology, actin structure, and cell adjacency that we had not yet utilized. Based on previous work that found a spatial correlation in the proliferation of PAEC, work showing that morphology can be used as a predictor of cellular response to molecular intervention, and recent work suggesting that genotypic and phenotypic traits are passed from mother to siblings cells, we hypothesized that we would find spatial clustering of cells with similar overall responses to IGF-1 administration.^5,6,35,38–41^

### 3.3 Spatial analysis of the single-cell response to IGF-1 administration

We next sought to build informative feature profiles of individual cells using the full set of measurements obtained from our imaging data (Figure 7). Using CellProfiler 3.1, we measured 390 features per cell.^33^ We normalized all measured features across experimental conditions, removed 1 uninformative feature, and calculated that 119 principal components accounted for 99% of the observed variance. Using this data, we asked *how likely are neighboring cells to be similar in any given condition?* The rationale for this question was our overall hypothesis that IGF-1 administration induced changes in specific sub-populations of PAEC. To test this hypothesis, we quantified the correlation of single-cell features with and without knowledge of the spatial position.

**Figure 7.**
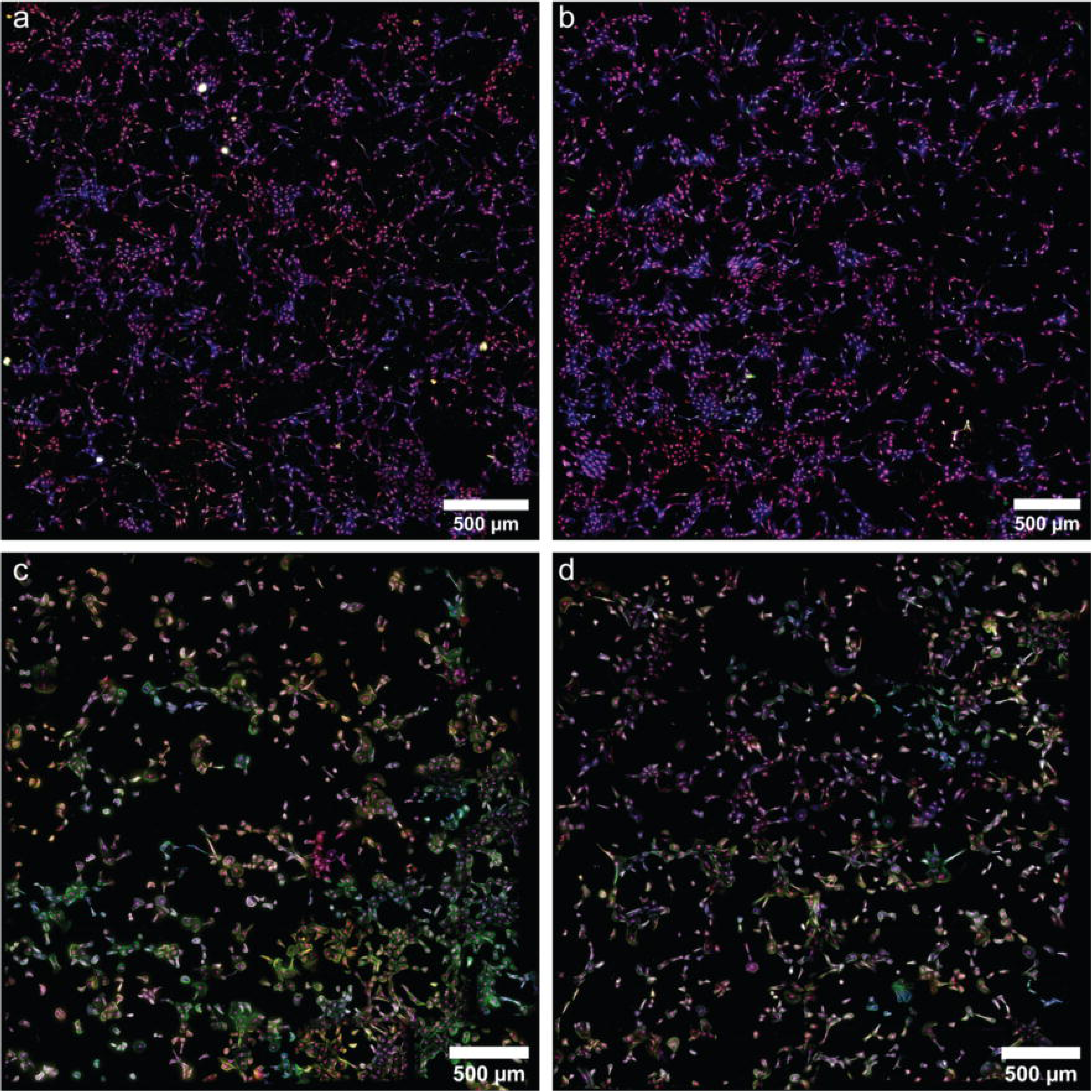
Representative high throughput imaging assays of nuclei (purple), actin (blue), Enos (green), and VEGF (red) in 18-hour tube formation assays for a) normal PAEC in vehicle media, b) normal PAEC in IGF-1 media, c) PPHN PAEC in vehicle media, d) PPHN PAEC in IGF-1 media.

By calculating the median Pearson correlation value for each cell’s nearest neighbors and comparing this to a null distribution of random sampling, we found that a distinct population of correlated cells emerge after IGF-1 administration in both normal and PPHN PAEC. This suggested there exist sub-populations of PAEC with a distinct response to IGF-1 treatment. To visualize this result, we recreated the spatial maps of cells but replaced each cell with a color code corresponding to uncorrelated cells (median Pearson correlation with nearest 20 cells less than the 95% value of the null distribution) and correlated cells (median Pearson correlation with nearest 20 cells greater than the 95% value of the null distribution) (Figure 8). This visualization confirmed that spatially correlated cells were clustered, were more likely at early time points after IGF-1 administration, and were more likely to occur in PPHN PAEC.

**Figure 8.**
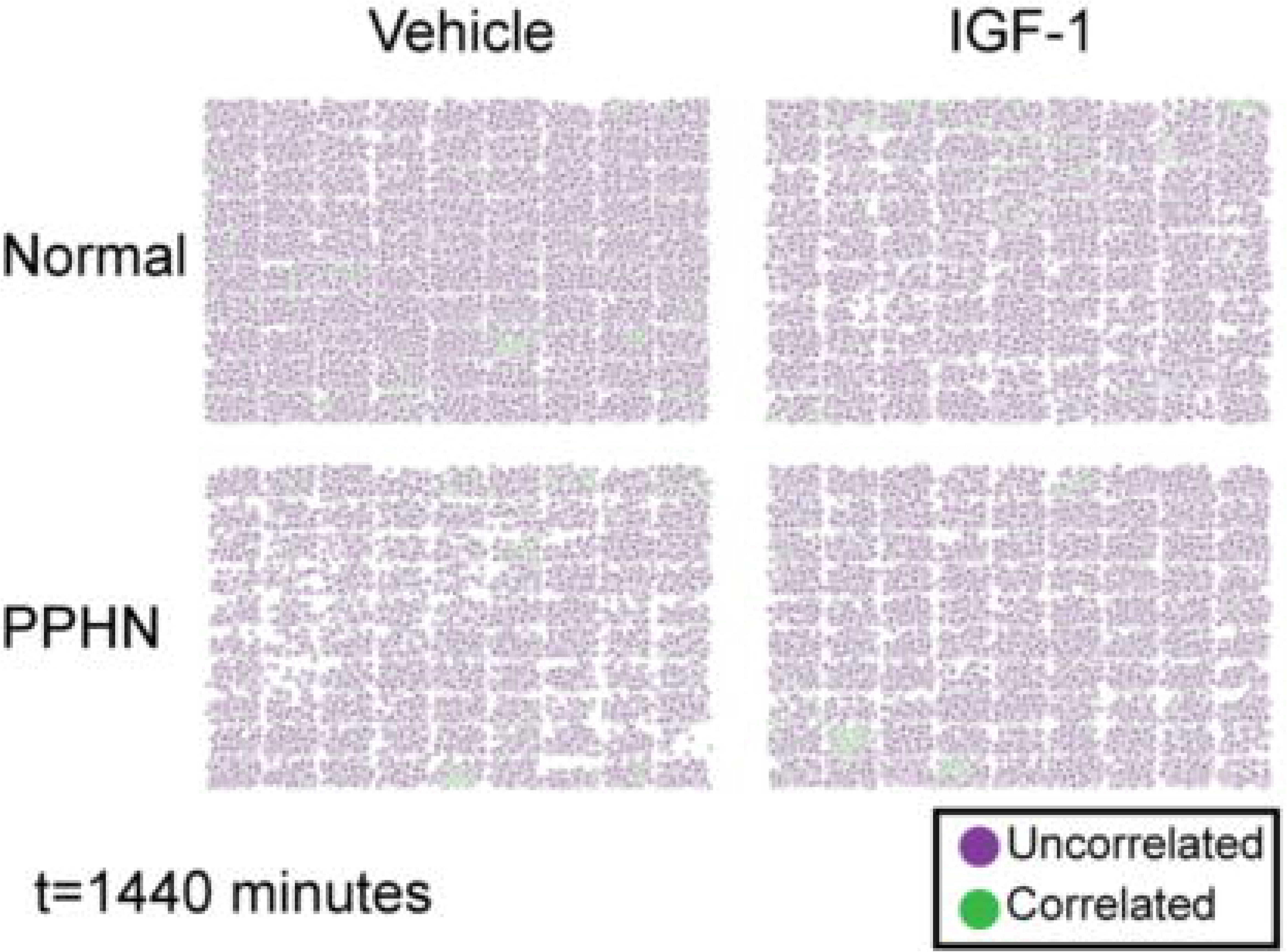
The spatial response of PAEC to IGF-1 at 1440 minutes. Purple cells are below the 95% non-spatial CDF value and are considered to be spatially uncorrelated to neighboring cells. Green cells are above the 95% non-spatial CDF value and considered to be spatially correlated to neighboring cells.

Finally, we calculated the percentage of PAEC exhibiting spatial correlations for each combination of time, media, and cell type (Figure 9). PPHN PAEC had a sustained increase in spatially correlated signaling after IGF-1 administration in our VEGF and eNOS imaging datasets. A similar sustained increase was not found in normal PAEC. Interestingly, the increase in spatial correlation after IGF-1 administration was transient for both normal and PPHN PAEC for the total protein imaging datasets. These findings strongly suggest that only a subset of spatially associated PPHN PAEC respond to IGF-1 administration. This IGF-1 responsive population may be responsible for the observed ensemble changes in proliferation, branch points, and diverse protein expression.

**Figure 9.**
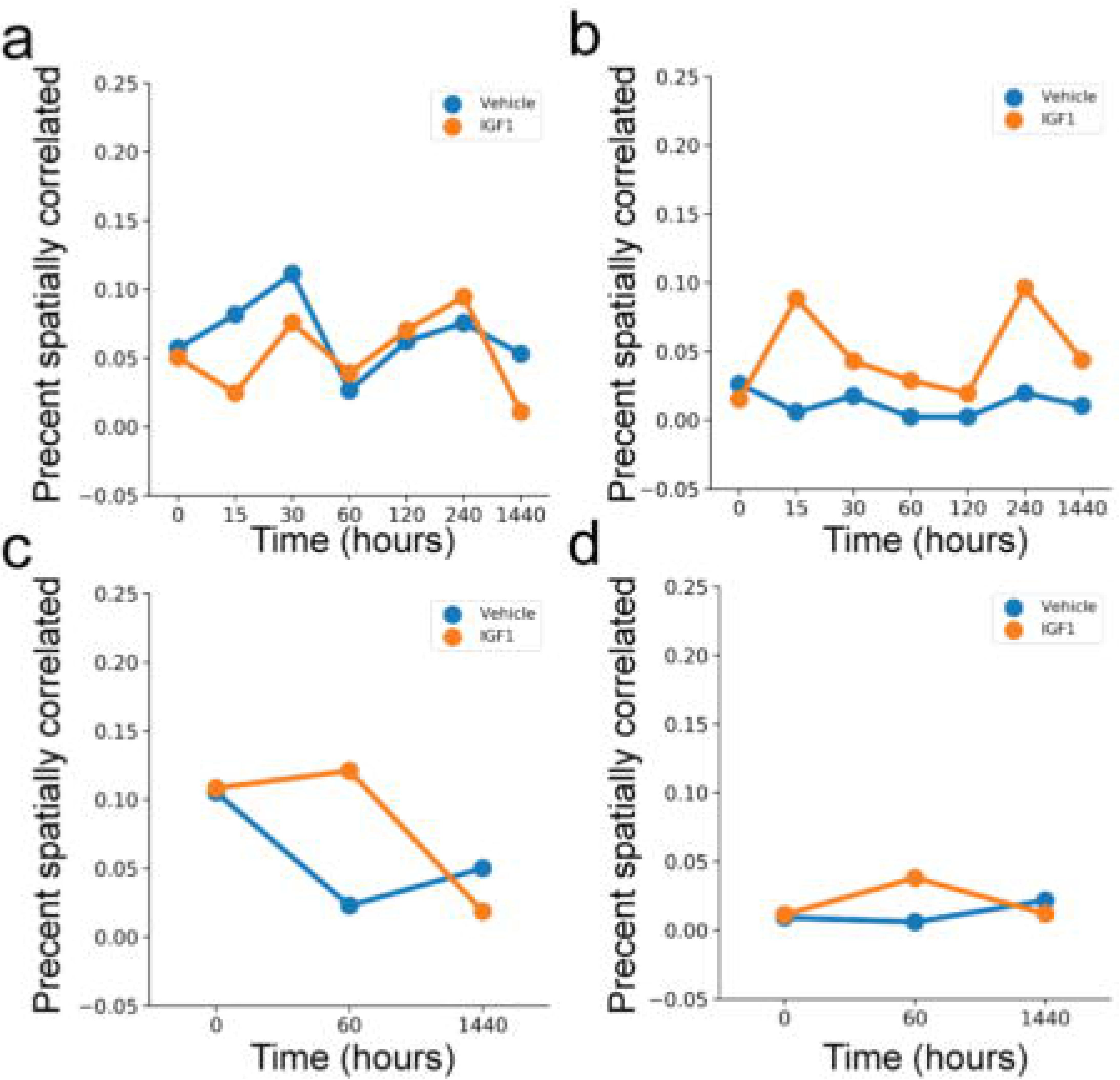
Percentage of spatially correlated PAEC in vehicle (blue) or IGF-1 (orange) media. Analysis of VEGF and eNOS immunofluorescence data for a) normal and b) PPHN PAEC. Analysis of total translation data for c) normal and d) PPHN PAEC.

## 4. Conclusion

In this study, we utilized a combination of traditional assays, high-throughput single-cell imaging, and statistical analyses to test if IGF-1 administration can improve the function of primary pulmonary artery endothelial cells. Using traditional biochemical assays, we found that IGF-1 increased proliferation and tube formation, but not angiogenic signaling, in PAEC isolated from healthy and PPHN sheep models. Previous studies have shown that regulation of the potent angiogenic molecules VEGF and eNOS is critical for the proper angiogenic function of PAEC.^25^

In primary sheep fetal PAEC, it is possible that IGF-1 does not act by regulating angiogenic signaling to increase proliferation and tube formation. This apparent conflict between protein expression and increased endothelial function led us to hypothesize there exist independent PAEC sub-populations with a varying response to IGF-1 administration. Motivated by our previous success on creating predictive models of changes to distributions in single-cell response to exogenous signaling using imaging-based assays, we utilized high-throughput single-cell imaging to image more than 1 million cells in systematic snapshots of cell type, treatment type, and time points following IGF-1 administration.^15,37^ Analyzing the obtained single-cell distributions, we confirmed that there exists a diverse PAEC response to IGF-1 administration.

Further analysis of these imaging data found that there is a sustained *in vitro* spatial dependence in PPHN PAEC response to IGF-1 administration. This finding suggests that beyond the observable heterogeneity, there are hidden variables, that we did not or cannot measure, contributing to PAEC regulation and response to IGF-1. One possible explanation for these observations is that each spatial cluster is the result of daughter cells from an initial PAEC that genetically encodes similar response to IGF-1.^5,6^ Another explanation is that certain PAEC excretes signaling molecules in response to IGF-1, creating localized signaling clusters.^42–45^ These two proposals are not mutually exclusive nor exhaustive. A key point to consider is that most previous work, including our own, on modeling single-cell heterogeneity assumes that each cell acts an independent unit for mathematical convenience.^15,37^ Integrating cell-to-cell communications is an exciting future direction. The high-density imaging dataset and pipeline made available through this work will help guide further studies.

Taken together, these results demonstrate that IGF-1 treatment partially restores normal endothelial cell behavior in primary PPHN PAEC. However, the signaling networks regulating PAEC response to IGF-1 treatment are not immediately clear from our experiments. What is clear is that there exist multiple sub-populations and that careful live-cell studies combined with high-throughput transcriptomic studies will be necessary to dissect the molecular pathways involved in IGF-1 response. This multi-generational genetic heritability of variable response has previously been observed in clonal cell lines.^5,46^ Here, the use of primary cells complicates the analysis of the data but also is closer to the *in vivo* reality of the disease phenotype. Other recent work has demonstrated that transcriptome data may poorly predict cellular phenotype, arguing that comprehensive studies are necessary to understand cellular function.^47,48^

Beyond the immediate experiment, these results have fundamental implications for how drug discovery is performed using *in vitro* experiments related to disease therapy. Standard metrics, such as proliferation and population-level readouts of cellular function, assume that drug treatment acts uniformly. In this study, high-throughput single-cell studies show that our observed outcomes are due to cellular subpopulations, each with different response to drug administration. The spatial correlation observed in this study is only possible using imaging-based assays that quantify both molecular readouts, cell morphology, and spatial positions. We propose that careful integration of non-spatial and spatial single cell studies are critical to fully understanding how potential molecular therapies interact with cell populations.

## Acknowledgments

The authors acknowledge Dr. Guy Hagen for assistance with using SIMToolbox to create the DMD to camera calibration mapping as well as Dr. Mark Dell’Acqua and Dr. Andrew Thorburn for insightful feedback. CK acknowledges funding from the University of Colorado Anschutz Medical Campus surgery resident training program. CK, GJS, SHA, and DPS acknowledge funding and IGF-1/BP3 reagent from Shire Pharmaceutics. GJS, SHA, and DPS acknowledge funding from NIH NHLBI 2R56HL068702-13. DPS acknowledges startup funding from the School of Medicine, University of Colorado Anschutz Medical Campus.

## Author contributions

SHA performed fetal sheep surgeries. GJS isolated and verified sheep PAEC. CK and GJS performed and analyzed PAEC culture, proliferation, and tube formation assays. CK and GJS cultured and labeled PAEC for high-throughput immunofluorescence imaging. CK, GJS, and DPS performed high-throughput immunofluorescence imaging. DPS designed and constructed the high-throughput fluorescence microscope. DPS designed and executed image analysis pipeline. CK and DPS performed image analysis. DPS designed and performed non-spatial and spatial single-cell analysis. SHA, GJS, and DPS designed the study. All authors wrote the manuscript.

## Supplemental Materials Description

Supplemental_figures.pdf - Supplemental figures 1-5

preprocess_raw_data.ijm - ImageJ script used to convert raw data to photons counts and arrange fluorescence channels for batch deconvolution.

process_deconvolved_data.ijm - ImageJ Script used to flat-field and maximum project all deconvolved images analyze_IF_data.cp - CellProfiler 3.1 pipeline used to quantify all VEGF/eNOS immunofluorescence data. The thresholding and filtering steps for the primary objects in the pipeline were manually checked for each imaging run.

analyze_Translation_data.cp - CellProfiler 3.1 pipeline used to quantify all translation data. The thresholding and filtering steps for the primary objects in the pipeline were manually checked for each imaging run. generate_figures.ipynb *-* Jupyter notebook to perform analysis and generate Figure 3-8 generate_supplemental_figures.ipynb - Jupyter notebook to generate Supplemental Figures 2-4.

