## Supplemental Figures for "High throughput imaging identifies a spatially localized response of primary fetal pulmonary artery endothelial cells to insulin-like growth factor 1 treatment"

### Supplemental Materials

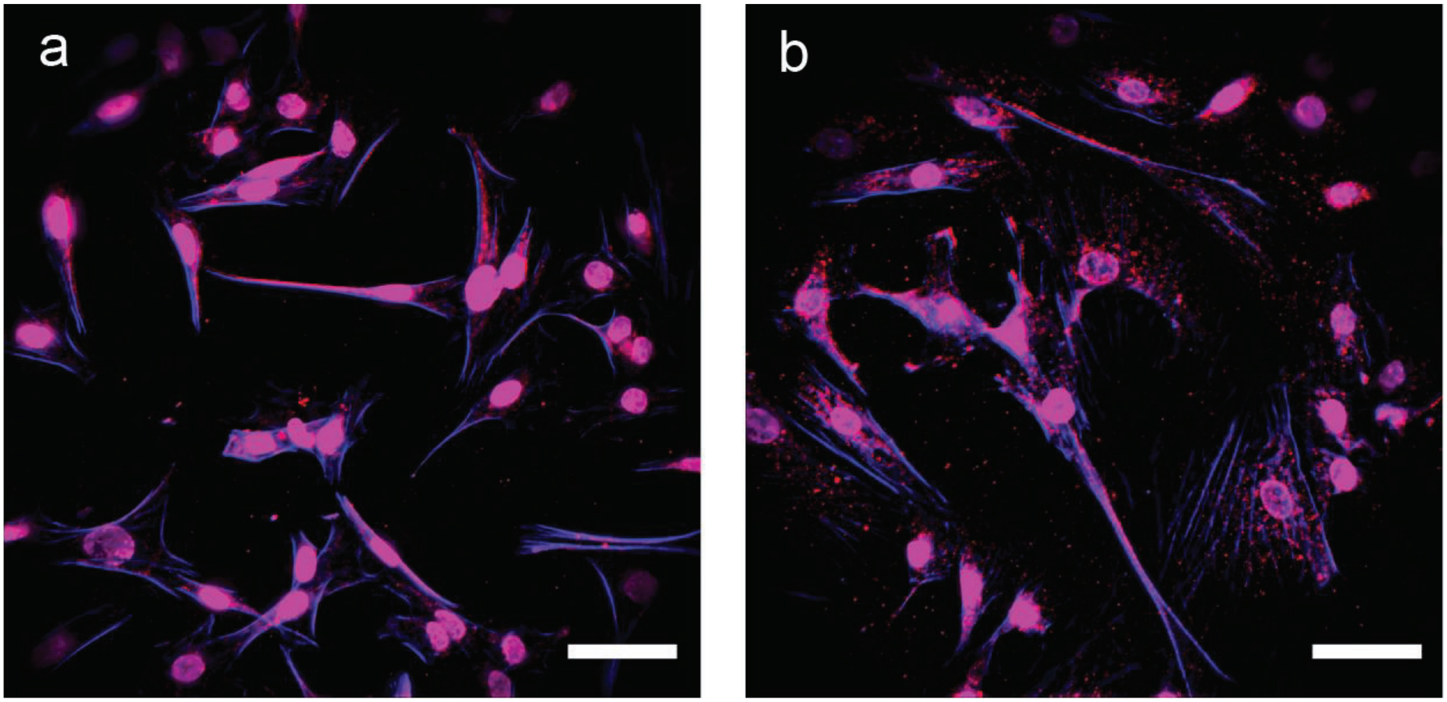


**Supplemental Figure 1.** vWF expression for a) normal and b) PPHN PAEC. (scale bar - 50 um)

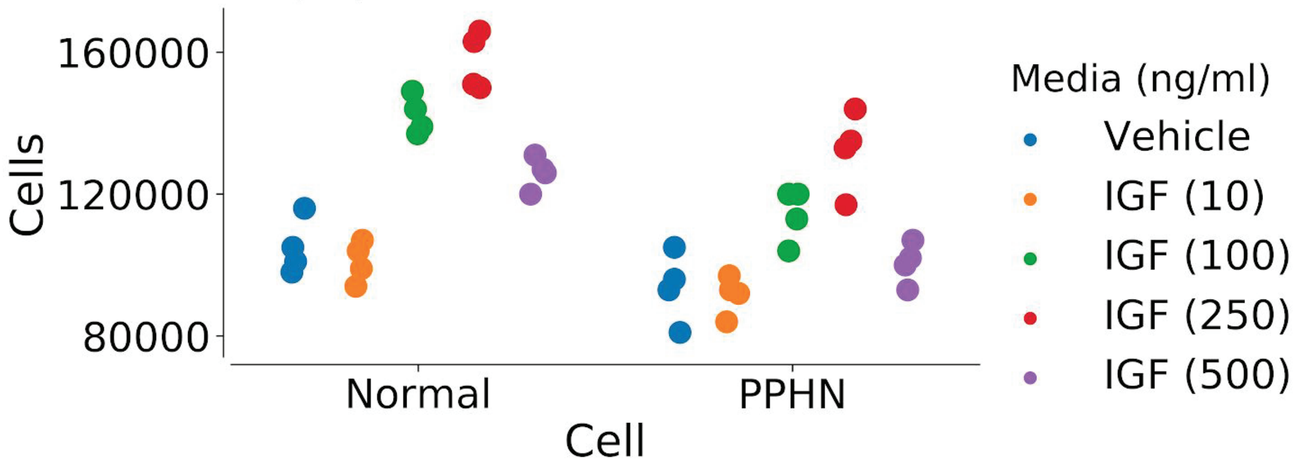


**Supplemental Figure 2.** IGF-1 dose response for normal and PPHN PAEC.

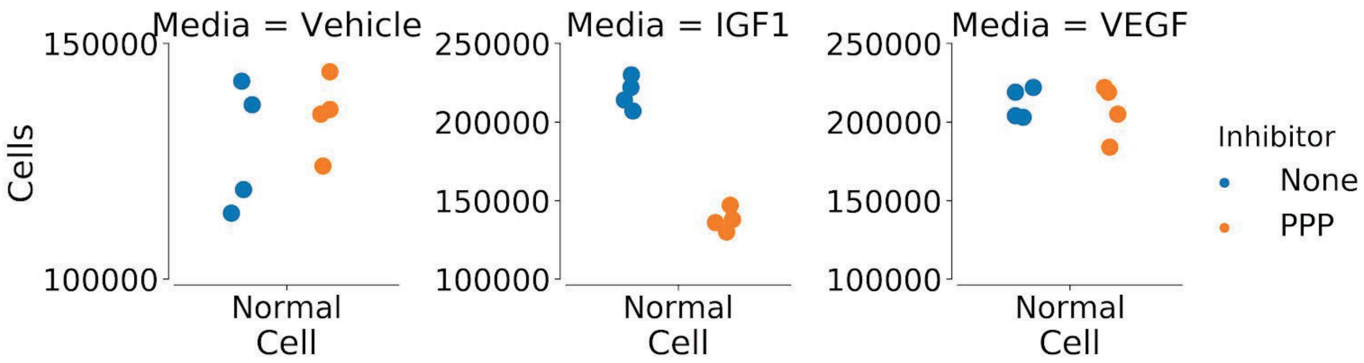


**Supplemental Figure 3.** Normal PAEC growth in the presence of 25 nM picropodophyllin (PPP) in vehicle, IGF-1 (250 ng/ml), and VEGF (50 ng/ml) media.

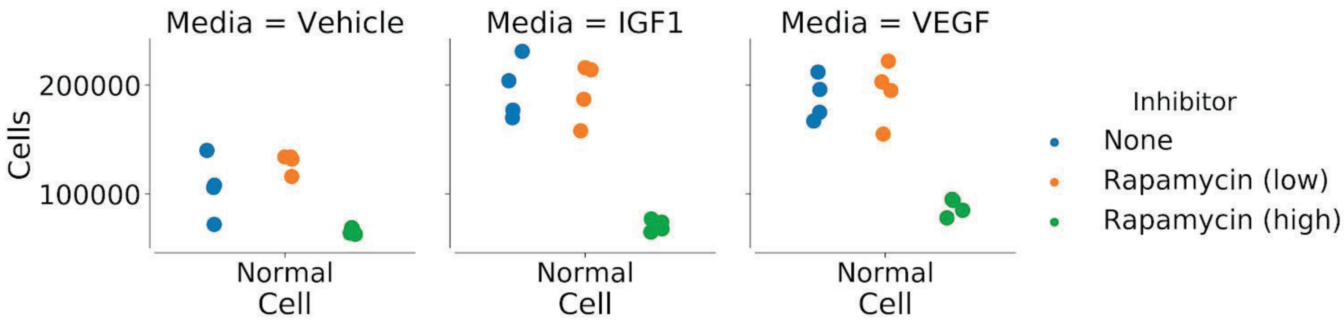


**Supplemental Figure 4.** Normal PAEC growth in the presence of low (.01 nM) and high (.1 nM) rapamycin (RP) in vehicle, IGF-1 (250 ng/ml), and VEGF (50 ng/ml) media.

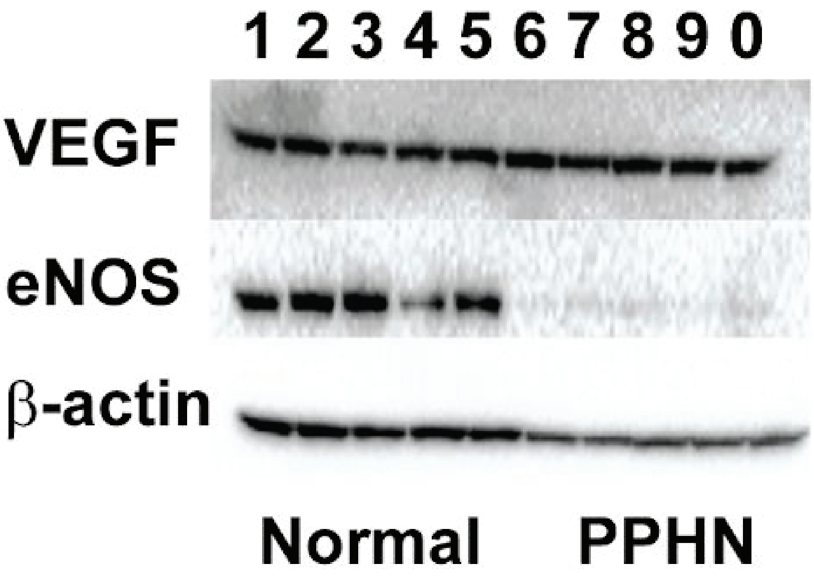


**Supplemental Figure 5.** One of three western blot experiments for data in Figure 3b. Key: (1 – 0 hour, normal PAEC; 2 – 1 hour, normal PAEC, vehicle media; 3 – 1 hour, normal PAEC, IGF-1 media; 4 – 24 hour, normal PAEC, vehicle media; 5 – 24 hour, normal PAEC, IGF-1 media; 6 – 0 hour, PPHN PAEC; 7 – 1 hour, PPHN PAEC, vehicle media; 8 – 1 hour, PPHN PAEC, IGF-1 media; 9 – 24 hour, PPHN PAEC, vehicle media; 0 – 24 hour, PPHN PAEC, IGF-1 media. There were no significant deviations observed in the two replicates of this experiment.
