## Supplemental code for "High throughput imaging identifies a spatially localized response of primary fetal pulmonary artery endothelial cells to insulin-like growth factor 1 treatment": generate_figures.ipynb

```
# %% [code]
# import libraries for analysis and plotting
import pandas as pd
import seaborn as sns
import numpy as np
import scipy as sp
import random
import matplotlib.pyplot as plt
from sklearn.preprocessing import StandardScaler
from sklearn.decomposition import PCA
from statsmodels import robust
import warnings
warnings.filterwarnings('ignore')

# This library must be downloaded and installed into the active environment 
# from: https://github.com/webermarcolivier/statannot
from statannot import add_stat_annotation 

# %% [code]
# Load ensemble data  

# Import western blot data
Ensemble_western_blots=pd.read_csv("ensemble/westernblot.csv")
Ensemble_western_blots['Type']='WB'

# Change Value to Normalized Intensity
Ensemble_western_blots.rename(index=str, columns={"Value": "Normalized Intensity"}, inplace=True)

# Create individual dataframes for statistical testing
Ensemble_western_blots[(Ensemble_western_blots.Cell == 'Normal') | (Ensemble_western_blots.Protein == 'VEGF')]
Ensemble_WB_Normal_VEGF=Ensemble_western_blots[(Ensemble_western_blots.Cell == 'Normal') | (Ensemble_western_blots.Protein == 'VEGF')]
Ensemble_WB_Normal_ENOS=Ensemble_western_blots[(Ensemble_western_blots.Cell == 'Normal') | (Ensemble_western_blots.Protein == 'eNOS')]
Ensemble_WB_PPHN_VEGF=Ensemble_western_blots[(Ensemble_western_blots.Cell == 'PPHN') | (Ensemble_western_blots.Protein == 'VEGF')]
Ensemble_WB_PPHN_ENOS=Ensemble_western_blots[(Ensemble_western_blots.Cell == 'PPHN') | (Ensemble_western_blots.Protein == 'eNOS')]

# Import proliferation data
Ensemble_proliferation=pd.read_csv("ensemble/proliferation.csv")
Ensemble_proliferation['Type']='Proliferation'

# Create individual data structures for statistical tests
Ensemble_proliferation_Normal=Ensemble_proliferation[Ensemble_proliferation['Cell'] == 'Normal']
Ensemble_proliferation_PPHN=Ensemble_proliferation[Ensemble_proliferation['Cell'] == 'PPHN']

# Import tube formation assays
Ensemble_tube_formation_assay=pd.read_csv("ensemble/tubeformation.csv")
Ensemble_tube_formation_assay['Type']='TFA'

# Create individual data structures for statistical tests
Ensemble_TFA_Normal=Ensemble_tube_formation_assay[Ensemble_tube_formation_assay['Cell'] == 'Normal']
Ensemble_TFA_PPHN=Ensemble_tube_formation_assay[Ensemble_tube_formation_assay['Cell'] == 'PPHN']

# Create merged data structures for nice plotting
merged_outer_1 = pd.merge(left=Ensemble_western_blots,right=Ensemble_proliferation, how='outer')
Ensemble_merged=pd.merge(left=merged_outer_1,right=Ensemble_tube_formation_assay, how='outer')

# %% [code]
#-------------------------------------------------------------------------------
# Plot and run statistical tests for ensemble data
# This generates the panels for Figures 3
#-------------------------------------------------------------------------------

# Run statistical tests and generate plots for western blot data

ax_western_blot = sns.catplot("Time (hours)", "Normalized Intensity", "Media", col="Cell", row="Protein",
                    data=Ensemble_western_blots, kind="strip", dodge=True, s=15, sharey=True)
sns.set_context("paper", font_scale=2)
axes_western_blot=ax_western_blot.axes

'''
add_stat_annotation(axes_western_blot[0,0], data=Ensemble_WB_Normal_VEGF, 
                    x="Time (hours)", y="Normalized Intensity", hue="Media",
                    boxPairList=[((1, "Vehicle"), (1, "IGF1")),
                                ((24, "Vehicle"), (24, "IGF1")),
                                ((0, "Vehicle"), (24, "Vehicle")),
                                ((0, "Vehicle"), (24, "IGF1"))],
                    test='Mann-Whitney', textFormat='star', loc='inside', verbose=2)

add_stat_annotation(axes_western_blot[0,1], data=Ensemble_WB_Normal_ENOS, 
                    x="Time (hours)", y="Normalized Intensity", hue="Media",
                    boxPairList=[((1, "Vehicle"), (1, "IGF1")),
                                ((24, "Vehicle"), (24, "IGF1")),
                                ((0, "Vehicle"), (24, "Vehicle")),
                                ((0, "Vehicle"), (24, "IGF1"))],
                    test='Mann-Whitney', textFormat='star', loc='inside', verbose=2)

add_stat_annotation(axes_western_blot[1,0], data=Ensemble_WB_PPHN_VEGF, 
                    x="Time (hours)", y="Normalized Intensity", hue="Media",
                    boxPairList=[((1, "Vehicle"), (1, "IGF1")),
                                ((24, "Vehicle"), (24, "IGF1")),
                                ((0, "Vehicle"), (24, "Vehicle")),
                                ((0, "Vehicle"), (24, "IGF1"))],
                    test='Mann-Whitney', textFormat='star', loc='inside', verbose=2)

add_stat_annotation(axes_western_blot[1,1], data=Ensemble_WB_PPHN_ENOS, 
                    x="Time (hours)", y="Normalized Intensity", hue="Media",
                    boxPairList=[((1, "Vehicle"), (1, "IGF1")),
                                ((24, "Vehicle"), (24, "IGF1")),
                                ((0, "Vehicle"), (24, "Vehicle")),
                                ((0, "Vehicle"), (24, "IGF1"))],
                    test='Mann-Whitney', textFormat='star', loc='inside', verbose=2)

axes_western_blot[0,0].set_yticks([0, 0.5, 1.0])
axes_western_blot[0,1].set_yticks([0, 0.5, 1.0])
axes_western_blot[1,0].set_yticks([0, 0.5, 1.0])
axes_western_blot[1,1].set_yticks([0, 0.5, 1.0])\
'''
sns.despine(offset=5, trim=False);

fig = ax_western_blot.fig
fig.savefig('figure_03/ensemble_western_blot.png', dpi=600)

# Run statistical tests and generate plots for proliferation data

ax_proliferation = sns.catplot("Day", "Cells", "Media", col="Cell", 
                               data=Ensemble_proliferation, kind="strip", dodge=True, s=15, sharey=True)
sns.set_context("paper", font_scale=2)
axes_proliferation=ax_proliferation.axes
'''
add_stat_annotation(axes_proliferation[0,0], data=Ensemble_proliferation_Normal, x="Day", y="Cells", hue="Media",
                    boxPairList=[((1, "Vehicle"), (1, "IGF1")),
                                ((2, "Vehicle"), (2, "IGF1")),
                                ((3, "Vehicle"), (3, "IGF1")),
                                ((0, "Vehicle"), (3, "Vehicle")),
                                ((0, "Vehicle"), (3, "IGF1"))],
                    test='Mann-Whitney', textFormat='star', loc='inside', verbose=2)

add_stat_annotation(axes_proliferation[0,1], data=Ensemble_proliferation_PPHN, x="Day", y="Cells", hue="Media",
                    boxPairList=[((1, "Vehicle"), (1, "IGF1")),
                                ((2, "Vehicle"), (2, "IGF1")),
                                ((3, "Vehicle"), (3, "IGF1")),
                                ((0, "Vehicle"), (3, "Vehicle")),
                                ((0, "Vehicle"), (3, "IGF1"))],
                    test='Mann-Whitney', textFormat='star', loc='inside', verbose=2)


axes_proliferation[0,0].set_yticks([300000, 600000])
axes_proliferation[0,1].set_yticks([100000, 200000])
'''
sns.despine(offset=5, trim=False);

fig = ax_proliferation.fig
fig.savefig('figure_03/ensemble_proliferation.png', dpi=600)

# Run statistical tests and generate plots for tube formation assay data

ax_tfa = sns.catplot( "Media","Branch points", col="Cell", data=Ensemble_tube_formation_assay, 
                     kind="strip", dodge=True, s=15, sharey=True)
sns.set_context("paper", font_scale=2)
axes_tfa=ax_tfa.axes
'''
add_stat_annotation(axes_tfa[0,0], data=Ensemble_tube_formation_assay, x="Cell", y="Branch points", hue="Media",
                    boxPairList=[(('Normal', "Vehicle"), ('Normal', "IGF1")),
                                (('PPHN', "Vehicle"), ('PPHN', "IGF1"))],
                    test='Mann-Whitney', textFormat='star', loc='inside', verbose=2)

axes_tfa[0,0].set_yticks([50, 100, 150, 200])
'''
sns.despine(offset=5, trim=False);
fig = ax_tfa.fig
fig.savefig('figure_03/ensemble_tube_formation_assay.png', dpi=600) 

# %% [code]
# Load single cell data

#-------------------------------------------------------------------------------
# Load vegf/enos IF assay data
#-------------------------------------------------------------------------------

# Load normal vehicle vegf/enos data
IF_Normal_VEH_time0000=pd.read_csv('IF/Normal/Vehicle/t0000/CP_RelabeledCellBody.txt', sep='\t')
IF_Normal_VEH_time0000['Cell']='Normal'
IF_Normal_VEH_time0000['Time']=0
IF_Normal_VEH_time0000['Media']='Vehicle'
IF_Normal_VEH_time0015=pd.read_csv('IF/Normal/Vehicle/t0015/CP_RelabeledCellBody.txt', sep='\t')
IF_Normal_VEH_time0015['Cell']='Normal'
IF_Normal_VEH_time0015['Time']=15
IF_Normal_VEH_time0015['Media']='Vehicle'
IF_Normal_VEH_time0030=pd.read_csv('IF/Normal/Vehicle/t0030/CP_RelabeledCellBody.txt', sep='\t')
IF_Normal_VEH_time0030['Cell']='Normal'
IF_Normal_VEH_time0030['Time']=30
IF_Normal_VEH_time0030['Media']='Vehicle'
IF_Normal_VEH_time0060=pd.read_csv('IF/Normal/Vehicle/t0060/CP_RelabeledCellBody.txt', sep='\t')
IF_Normal_VEH_time0060['Cell']='Normal'
IF_Normal_VEH_time0060['Time']=60
IF_Normal_VEH_time0060['Media']='Vehicle'
IF_Normal_VEH_time0120=pd.read_csv('IF/Normal/Vehicle/t0120/CP_RelabeledCellBody.txt', sep='\t')
IF_Normal_VEH_time0120['Cell']='Normal'
IF_Normal_VEH_time0120['Time']=120
IF_Normal_VEH_time0120['Media']='Vehicle'
IF_Normal_VEH_time0240=pd.read_csv('IF/Normal/Vehicle/t0240/CP_RelabeledCellBody.txt', sep='\t')
IF_Normal_VEH_time0240['Cell']='Normal'
IF_Normal_VEH_time0240['Time']=240
IF_Normal_VEH_time0240['Media']='Vehicle'
IF_Normal_VEH_time1440=pd.read_csv('IF/Normal/Vehicle/t1440/CP_RelabeledCellBody.txt', sep='\t')
IF_Normal_VEH_time1440['Cell']='Normal'
IF_Normal_VEH_time1440['Time']=1440
IF_Normal_VEH_time1440['Media']='Vehicle'

# Create normal vehicle vegf/enos data structure
IF_Normal_VEH=pd.concat((IF_Normal_VEH_time0000,IF_Normal_VEH_time0015,IF_Normal_VEH_time0030,
                 IF_Normal_VEH_time0060,IF_Normal_VEH_time0120,IF_Normal_VEH_time0240,
                     IF_Normal_VEH_time1440),axis=0,sort=False)

# Load normal IGF1 vegf/enos data
IF_Normal_IGF_time0000=pd.read_csv('IF/Normal/IGF1/t0000/CP_RelabeledCellBody.txt', sep='\t')
IF_Normal_IGF_time0000['Cell']='Normal'
IF_Normal_IGF_time0000['Time']=0
IF_Normal_IGF_time0000['Media']='IGF1'
IF_Normal_IGF_time0015=pd.read_csv('IF/Normal/IGF1/t0015/CP_RelabeledCellBody.txt', sep='\t')
IF_Normal_IGF_time0015['Cell']='Normal'
IF_Normal_IGF_time0015['Time']=15
IF_Normal_IGF_time0015['Media']='IGF1'
IF_Normal_IGF_time0030=pd.read_csv('IF/Normal/IGF1/t0030/CP_RelabeledCellBody.txt', sep='\t')
IF_Normal_IGF_time0030['Cell']='Normal'
IF_Normal_IGF_time0030['Time']=30
IF_Normal_IGF_time0030['Media']='IGF1'
IF_Normal_IGF_time0060=pd.read_csv('IF/Normal/IGF1/t0060/CP_RelabeledCellBody.txt', sep='\t')
IF_Normal_IGF_time0060['Cell']='Normal'
IF_Normal_IGF_time0060['Time']=60
IF_Normal_IGF_time0060['Media']='IGF1'
IF_Normal_IGF_time0120=pd.read_csv('IF/Normal/IGF1/t0120/CP_RelabeledCellBody.txt', sep='\t')
IF_Normal_IGF_time0120['Cell']='Normal'
IF_Normal_IGF_time0120['Time']=120
IF_Normal_IGF_time0120['Media']='IGF1'
IF_Normal_IGF_time0240=pd.read_csv('IF/Normal/IGF1/t0240/CP_RelabeledCellBody.txt', sep='\t')
IF_Normal_IGF_time0240['Cell']='Normal'
IF_Normal_IGF_time0240['Time']=240
IF_Normal_IGF_time0240['Media']='IGF1'
IF_Normal_IGF_time1440=pd.read_csv('IF/Normal/IGF1/t1440/CP_RelabeledCellBody.txt', sep='\t')
IF_Normal_IGF_time1440['Cell']='Normal'
IF_Normal_IGF_time1440['Time']=1440
IF_Normal_IGF_time1440['Media']='IGF1'

# Create normal IGF1 vegf/enos data structure
IF_Normal_IGF=pd.concat((IF_Normal_IGF_time0000,IF_Normal_IGF_time0015,IF_Normal_IGF_time0030,
                     IF_Normal_IGF_time0060,IF_Normal_IGF_time0120,IF_Normal_IGF_time0240,
                     IF_Normal_IGF_time1440),axis=0,sort=False)

# Load PPHN vehicle vegf/enos data
IF_PPHN_VEH_time0000=pd.read_csv('IF/PPHN/Vehicle/t0000/CP_RelabeledCellBody.txt', sep='\t')
IF_PPHN_VEH_time0000['Cell']='PPHN'
IF_PPHN_VEH_time0000['Time']=0
IF_PPHN_VEH_time0000['Media']='Vehicle'
IF_PPHN_VEH_time0015=pd.read_csv('IF/PPHN/Vehicle/t0015/CP_RelabeledCellBody.txt', sep='\t')
IF_PPHN_VEH_time0015['Cell']='PPHN'
IF_PPHN_VEH_time0015['Time']=15
IF_PPHN_VEH_time0015['Media']='Vehicle'
IF_PPHN_VEH_time0030=pd.read_csv('IF/PPHN/Vehicle/t0030/CP_RelabeledCellBody.txt', sep='\t')
IF_PPHN_VEH_time0030['Cell']='PPHN'
IF_PPHN_VEH_time0030['Time']=30
IF_PPHN_VEH_time0030['Media']='Vehicle'
IF_PPHN_VEH_time0060=pd.read_csv('IF/PPHN/Vehicle/t0060/CP_RelabeledCellBody.txt', sep='\t')
IF_PPHN_VEH_time0060['Cell']='PPHN'
IF_PPHN_VEH_time0060['Time']=60
IF_PPHN_VEH_time0060['Media']='Vehicle'
IF_PPHN_VEH_time0120=pd.read_csv('IF/PPHN/Vehicle/t0120/CP_RelabeledCellBody.txt', sep='\t')
IF_PPHN_VEH_time0120['Cell']='PPHN'
IF_PPHN_VEH_time0120['Time']=120
IF_PPHN_VEH_time0120['Media']='Vehicle'
IF_PPHN_VEH_time0240=pd.read_csv('IF/PPHN/Vehicle/t0240/CP_RelabeledCellBody.txt', sep='\t')
IF_PPHN_VEH_time0240['Cell']='PPHN'
IF_PPHN_VEH_time0240['Time']=240
IF_PPHN_VEH_time0240['Media']='Vehicle'
IF_PPHN_VEH_time1440=pd.read_csv('IF/PPHN/Vehicle/t1440/CP_RelabeledCellBody.txt', sep='\t')
IF_PPHN_VEH_time1440['Cell']='PPHN'
IF_PPHN_VEH_time1440['Time']=1440
IF_PPHN_VEH_time1440['Media']='Vehicle'

# Create PPHN vehicle vegf/enos data structure
IF_PPHN_VEH=pd.concat((IF_PPHN_VEH_time0000,IF_PPHN_VEH_time0015,IF_PPHN_VEH_time0030,
                     IF_PPHN_VEH_time0060,IF_PPHN_VEH_time0120,IF_PPHN_VEH_time0240,
                     IF_PPHN_VEH_time1440),axis=0,sort=False)

# Load PPHN IGF1 vegf/enos data
IF_PPHN_IGF_time0000=pd.read_csv('IF/PPHN/IGF1/t0000/CP_RelabeledCellBody.txt', sep='\t')
IF_PPHN_IGF_time0000['Cell']='PPHN'
IF_PPHN_IGF_time0000['Time']=0
IF_PPHN_IGF_time0000['Media']='IGF1'
IF_PPHN_IGF_time0015=pd.read_csv('IF/PPHN/IGF1/t0015/CP_RelabeledCellBody.txt', sep='\t')
IF_PPHN_IGF_time0015['Cell']='PPHN'
IF_PPHN_IGF_time0015['Time']=15
IF_PPHN_IGF_time0015['Media']='IGF1'
IF_PPHN_IGF_time0030=pd.read_csv('IF/PPHN/IGF1/t0030/CP_RelabeledCellBody.txt', sep='\t')
IF_PPHN_IGF_time0030['Cell']='PPHN'
IF_PPHN_IGF_time0030['Time']=30
IF_PPHN_IGF_time0030['Media']='IGF1'
IF_PPHN_IGF_time0060=pd.read_csv('IF/PPHN/IGF1/t0060/CP_RelabeledCellBody.txt', sep='\t')
IF_PPHN_IGF_time0060['Cell']='PPHN'
IF_PPHN_IGF_time0060['Time']=60
IF_PPHN_IGF_time0060['Media']='IGF1'
IF_PPHN_IGF_time0120=pd.read_csv('IF/PPHN/IGF1/t0120/CP_RelabeledCellBody.txt', sep='\t')
IF_PPHN_IGF_time0120['Cell']='PPHN'
IF_PPHN_IGF_time0120['Time']=120
IF_PPHN_IGF_time0120['Media']='IGF1'
IF_PPHN_IGF_time0240=pd.read_csv('IF/PPHN/IGF1/t0240/CP_RelabeledCellBody.txt', sep='\t')
IF_PPHN_IGF_time0240['Cell']='PPHN'
IF_PPHN_IGF_time0240['Time']=240
IF_PPHN_IGF_time0240['Media']='IGF1'
IF_PPHN_IGF_time1440=pd.read_csv('IF/PPHN/IGF1/t1440/CP_RelabeledCellBody.txt', sep='\t')
IF_PPHN_IGF_time1440['Cell']='PPHN'
IF_PPHN_IGF_time1440['Time']=1440
IF_PPHN_IGF_time1440['Media']='IGF1'

# Create PPHN IGF1 vegf/enos data structure
IF_PPHN_IGF=pd.concat((IF_PPHN_IGF_time0000,IF_PPHN_IGF_time0015,IF_PPHN_IGF_time0030,
                     IF_PPHN_IGF_time0060,IF_PPHN_IGF_time0120,IF_PPHN_IGF_time0240,
                     IF_PPHN_IGF_time1440),axis=0,sort=False)

#-------------------------------------------------------------------------------
# Load translation assay data
#-------------------------------------------------------------------------------

# Load normal vehicle translation data
Translation_Normal_VEH_time0000=pd.read_csv('Translation/Normal/Vehicle/t0000/CP_RelabeledCellBody.txt', sep='\t')
Translation_Normal_VEH_time0000['Cell']='Normal'
Translation_Normal_VEH_time0000['Time']=0
Translation_Normal_VEH_time0000['Media']='Vehicle'

Translation_Normal_VEH_time0060=pd.read_csv('Translation/Normal/Vehicle/t0060/CP_RelabeledCellBody.txt', sep='\t')
Translation_Normal_VEH_time0060['Cell']='Normal'
Translation_Normal_VEH_time0060['Time']=60
Translation_Normal_VEH_time0060['Media']='Vehicle'

Translation_Normal_VEH_time1440=pd.read_csv('Translation/Normal/Vehicle/t1440/CP_RelabeledCellBody.txt', sep='\t')
Translation_Normal_VEH_time1440['Cell']='Normal'
Translation_Normal_VEH_time1440['Time']=1440
Translation_Normal_VEH_time1440['Media']='Vehicle'

# Create normal vehicle translation data structure
Translation_Normal_VEH=pd.concat((Translation_Normal_VEH_time0000,Translation_Normal_VEH_time0060,Translation_Normal_VEH_time1440),axis=0,sort=False)

# Load normal IGF1 translation data
Translation_Normal_IGF_time0000=pd.read_csv('Translation/Normal/Vehicle/t0000/CP_RelabeledCellBody.txt', sep='\t')
Translation_Normal_IGF_time0000['Cell']='Normal'
Translation_Normal_IGF_time0000['Time']=0
Translation_Normal_IGF_time0000['Media']='IGF1'

Translation_Normal_IGF_time0060=pd.read_csv('Translation/Normal/IGF1/t0060/CP_RelabeledCellBody.txt', sep='\t')
Translation_Normal_IGF_time0060['Cell']='Normal'
Translation_Normal_IGF_time0060['Time']=60
Translation_Normal_IGF_time0060['Media']='IGF1'

Translation_Normal_IGF_time1440=pd.read_csv('Translation/Normal/IGF1/t1440/CP_RelabeledCellBody.txt', sep='\t')
Translation_Normal_IGF_time1440['Cell']='Normal'
Translation_Normal_IGF_time1440['Time']=1440
Translation_Normal_IGF_time1440['Media']='IGF1'

# Create normal IGF1 translation data structure
Translation_Normal_IGF=pd.concat((Translation_Normal_IGF_time0000,Translation_Normal_IGF_time0060,Translation_Normal_IGF_time1440),axis=0,sort=False)

# Load PPHN vehicle translation data
Translation_PPHN_VEH_time0000=pd.read_csv('Translation/PPHN/Vehicle/t0000/CP_RelabeledCellBody.txt', sep='\t')
Translation_PPHN_VEH_time0000['Cell']='PPHN'
Translation_PPHN_VEH_time0000['Time']=0
Translation_PPHN_VEH_time0000['Media']='Vehicle'

Translation_PPHN_VEH_time0060=pd.read_csv('Translation/PPHN/Vehicle/t0060/CP_RelabeledCellBody.txt', sep='\t')
Translation_PPHN_VEH_time0060['Cell']='PPHN'
Translation_PPHN_VEH_time0060['Time']=60
Translation_PPHN_VEH_time0060['Media']='Vehicle'

Translation_PPHN_VEH_time1440=pd.read_csv('Translation/PPHN/Vehicle/t1440/CP_RelabeledCellBody.txt', sep='\t')
Translation_PPHN_VEH_time1440['Cell']='PPHN'
Translation_PPHN_VEH_time1440['Time']=1440
Translation_PPHN_VEH_time1440['Media']='Vehicle'

# Create PPHN vehicle translation data structure
Translation_PPHN_VEH=pd.concat((Translation_PPHN_VEH_time0000,Translation_PPHN_VEH_time0060,Translation_PPHN_VEH_time1440),axis=0,sort=False)

# Load PPHN IGF1 translation data
Translation_PPHN_IGF_time0000 = pd.read_csv('Translation/PPHN/Vehicle/t0000/CP_RelabeledCellBody.txt', sep='\t')
Translation_PPHN_IGF_time0000['Cell'] = 'PPHN'
Translation_PPHN_IGF_time0000['Time'] = 0
Translation_PPHN_IGF_time0000['Media'] = 'IGF1'

Translation_PPHN_IGF_time0060 = pd.read_csv('Translation/PPHN/IGF1/t0060/CP_RelabeledCellBody.txt', sep='\t')
Translation_PPHN_IGF_time0060['Cell'] = 'PPHN'
Translation_PPHN_IGF_time0060['Time'] = 60
Translation_PPHN_IGF_time0060['Media'] = 'IGF1'

Translation_PPHN_IGF_time1440 = pd.read_csv('Translation/PPHN/IGF1/t1440/CP_RelabeledCellBody.txt', sep='\t')
Translation_PPHN_IGF_time1440['Cell'] = 'PPHN'
Translation_PPHN_IGF_time1440['Time'] = 1440
Translation_PPHN_IGF_time1440['Media'] = 'IGF1'

# Create PPHN IGF1 translation data structure
Translation_PPHN_IGF = pd.concat((Translation_PPHN_IGF_time0000,Translation_PPHN_IGF_time0060,Translation_PPHN_IGF_time1440),axis=0,sort=False)

# %% [code]
# Define columns that are uniformative to cell features
toDrop = ['AreaShape_Center_X', 'AreaShape_Center_Y', 'AreaShape_Center_Z', 
          'Time', 'Media', 'Cell','ImageNumber','ObjectNumber']

# Define metadata columns
metaDataCols = ['Cell', 'Media', 'Time', 'ImageNumber','ObjectNumber',
                 'AreaShape_Center_X','AreaShape_Center_Y']

# %% [code]
#-------------------------------------------------------------------------------
# Determine uniformative features for vegf/enos IF data
#-------------------------------------------------------------------------------

# Calculate MAD for each column for cells at t=0 for normal cells in vehicle media
IF_Normal_Baseline=IF_Normal_VEH_time0000.dropna(axis=0)
IF_Normal_Baseline.reset_index(drop=True, inplace=True)
IF_Normal_Baseline.drop(toDrop,axis=1,inplace=True)
IF_Normal_Baseline.reset_index(drop=True, inplace=True)
IF_Normal_cols = IF_Normal_Baseline.columns

IF_Normal_data = robust.mad(IF_Normal_Baseline)
IF_Normal_dfMAD = pd.DataFrame(columns=IF_Normal_cols,data=np.atleast_2d(IF_Normal_data))

# Find all columns with MAD = 0
IF_Normal_colsToDropMAD = IF_Normal_dfMAD.columns[IF_Normal_dfMAD.eq(0).any()]
IF_Normal_dfMAD.drop(IF_Normal_colsToDropMAD,axis=1,inplace=True)

# Calculate MAD for each column for cells at t=0 for PPHN cells in vehicle media
IF_PPHN_Baseline = IF_PPHN_VEH_time0000.dropna(axis=0)
IF_PPHN_Baseline.reset_index(drop=True, inplace=True)
IF_PPHN_Baseline.drop(toDrop,axis=1,inplace=True)
IF_PPHN_Baseline.reset_index(drop=True, inplace=True)
IF_PPHN_cols = IF_PPHN_Baseline.columns

IF_PPHN_data = robust.mad(IF_PPHN_Baseline)
IF_PPHN_dfMAD = pd.DataFrame(columns=IF_PPHN_cols,data=np.atleast_2d(IF_PPHN_data))

# Find all columns with MAD = 0
IF_PPHN_colsToDropMAD = IF_PPHN_dfMAD.columns[IF_PPHN_dfMAD.eq(0).any()]
IF_PPHN_dfMAD.drop(IF_PPHN_colsToDropMAD,axis=1,inplace=True)

#-------------------------------------------------------------------------------
# Determine uniformative features for translation data
#-------------------------------------------------------------------------------

Translation_Normal_Baseline=Translation_Normal_VEH_time0000.dropna(axis=0)
Translation_Normal_Baseline.reset_index(drop=True, inplace=True)
Translation_Normal_Baseline.drop(toDrop,axis=1,inplace=True)
Translation_Normal_Baseline.reset_index(drop=True, inplace=True)
Translation_Normal_cols = Translation_Normal_Baseline.columns

Translation_Normal_data = robust.mad(Translation_Normal_Baseline)
Translation_Normal_dfMAD = pd.DataFrame(columns=Translation_Normal_cols,data=np.atleast_2d(Translation_Normal_data))

# Find all columns with MAD = 0
Translation_Normal_colsToDropMAD = Translation_Normal_dfMAD.columns[Translation_Normal_dfMAD.eq(0).any()]
Translation_Normal_dfMAD.drop(Translation_Normal_colsToDropMAD,axis=1,inplace=True)

Translation_PPHN_Baseline=Translation_PPHN_VEH_time0000.dropna(axis=0)
Translation_PPHN_Baseline.reset_index(drop=True, inplace=True)
Translation_PPHN_Baseline.drop(toDrop,axis=1,inplace=True)
Translation_PPHN_Baseline.reset_index(drop=True, inplace=True)
Translation_PPHN_cols = Translation_PPHN_Baseline.columns

Translation_PPHN_data = robust.mad(Translation_PPHN_Baseline)
Translation_PPHN_dfMAD = pd.DataFrame(columns=Translation_PPHN_cols,data=np.atleast_2d(Translation_PPHN_data))

# Find all columns with MAD = 0
Translation_PPHN_colsToDropMAD = Translation_PPHN_dfMAD.columns[Translation_PPHN_dfMAD.eq(0).any()]
Translation_PPHN_dfMAD.drop(Translation_PPHN_colsToDropMAD,axis=1,inplace=True)

# %% [code]
# Find and remove uniformative features

#-------------------------------------------------------------------------------
# Drop uniformative features (MAD=0) and features with NaN values for vegf/enos IF data
#-------------------------------------------------------------------------------

# Extract metadata, drop any rows with NaN values, drop any columns with MAD=0, and extract informative columns for PCA 
IF_Normal_VEH.dropna(axis=0, inplace=True)
IF_Normal_VEH.reset_index(drop=True, inplace=True)
IF_Normal_VEH_metaData = IF_Normal_VEH[metaDataCols]
IF_Normal_VEH.drop(toDrop,axis=1,inplace=True)
IF_Normal_VEH.drop(IF_Normal_colsToDropMAD,axis=1,inplace=True)
IF_Normal_VEH.reset_index(drop=True, inplace=True)

IF_Normal_IGF.dropna(axis=0, inplace=True)
IF_Normal_IGF.reset_index(drop=True, inplace=True)
IF_Normal_IGF_metaData = IF_Normal_IGF[metaDataCols]
IF_Normal_IGF.drop(toDrop,axis=1,inplace=True)
IF_Normal_IGF.drop(IF_Normal_colsToDropMAD,axis=1,inplace=True)
IF_Normal_IGF.reset_index(drop=True, inplace=True)

# Extract metadata, drop any rows with NaN values, drop any columns with MAD=0, and extract informative columns for PCA 
IF_PPHN_VEH.dropna(axis=0, inplace=True)
IF_PPHN_VEH.reset_index(drop=True, inplace=True)
IF_PPHN_VEH_metaData = IF_PPHN_VEH[metaDataCols]
IF_PPHN_VEH.drop(toDrop,axis=1,inplace=True)
IF_PPHN_VEH.drop(IF_PPHN_colsToDropMAD,axis=1,inplace=True)
IF_PPHN_VEH.reset_index(drop=True, inplace=True)

IF_PPHN_IGF.dropna(axis=0, inplace=True)
IF_PPHN_IGF.reset_index(drop=True, inplace=True)
IF_PPHN_IGF_metaData = IF_PPHN_IGF[metaDataCols]
IF_PPHN_IGF.drop(toDrop,axis=1,inplace=True)
IF_PPHN_IGF.drop(IF_PPHN_colsToDropMAD,axis=1,inplace=True)
IF_PPHN_IGF.reset_index(drop=True, inplace=True)

#-------------------------------------------------------------------------------
# Drop uniformative features and features with NaN values for translation data
#-------------------------------------------------------------------------------

# Extract metadata, drop any rows with NaN values, drop any columns with MAD=0, and extract informative columns for PCA 
Translation_Normal_VEH.dropna(axis=0, inplace=True)
Translation_Normal_VEH.reset_index(drop=True, inplace=True)
Translation_Normal_VEH_metaData = Translation_Normal_VEH[metaDataCols]
Translation_Normal_VEH.drop(toDrop,axis=1,inplace=True)
Translation_Normal_VEH.drop(Translation_Normal_colsToDropMAD,axis=1,inplace=True)
Translation_Normal_VEH.reset_index(drop=True, inplace=True)

Translation_Normal_IGF.dropna(axis=0, inplace=True)
Translation_Normal_IGF.reset_index(drop=True, inplace=True)
Translation_Normal_IGF_metaData = Translation_Normal_IGF[metaDataCols]
Translation_Normal_IGF.drop(toDrop,axis=1,inplace=True)
Translation_Normal_IGF.drop(Translation_Normal_colsToDropMAD,axis=1,inplace=True)
Translation_Normal_IGF.reset_index(drop=True, inplace=True)

# Extract metadata, drop any rows with NaN values, drop any columns with MAD=0, and extract informative columns for PCA 
Translation_PPHN_VEH.dropna(axis=0, inplace=True)
Translation_PPHN_VEH.reset_index(drop=True, inplace=True)
Translation_PPHN_VEH_metaData = Translation_PPHN_VEH[metaDataCols]
Translation_PPHN_VEH.drop(toDrop,axis=1,inplace=True)
Translation_PPHN_VEH.drop(Translation_PPHN_colsToDropMAD,axis=1,inplace=True)
Translation_PPHN_VEH.reset_index(drop=True, inplace=True)

Translation_PPHN_IGF.dropna(axis=0, inplace=True)
Translation_PPHN_IGF.reset_index(drop=True, inplace=True)
Translation_PPHN_IGF_metaData = Translation_PPHN_IGF[metaDataCols]
Translation_PPHN_IGF.drop(toDrop,axis=1,inplace=True)
Translation_PPHN_IGF.drop(Translation_PPHN_colsToDropMAD,axis=1,inplace=True)
Translation_PPHN_IGF.reset_index(drop=True, inplace=True)

# %% [code]
# Combine assays into one large structure for each experiment and cell type

#-------------------------------------------------------------------------------
# Create one large data structure for each cell type in vegf/enos IF data
#-------------------------------------------------------------------------------

IF_Normal_fusedData = pd.concat((IF_Normal_VEH,IF_Normal_IGF))
IF_Normal_fusedData.reset_index(drop=True, inplace=True)

IF_Normal_fusedmetaData = pd.concat((IF_Normal_VEH_metaData,IF_Normal_IGF_metaData))
IF_Normal_fusedmetaData.reset_index(drop=True, inplace=True)
                                    
IF_PPHN_fusedData = pd.concat((IF_PPHN_VEH,IF_PPHN_IGF))
IF_PPHN_fusedData.reset_index(drop=True, inplace=True)

IF_PPHN_fusedmetaData = pd.concat((IF_PPHN_VEH_metaData,IF_PPHN_IGF_metaData))
IF_PPHN_fusedmetaData.reset_index(drop=True, inplace=True)
                                  
#-------------------------------------------------------------------------------
# Create one large data structure for each cell type in translation data
#-------------------------------------------------------------------------------

Translation_Normal_fusedData = pd.concat((Translation_Normal_VEH,Translation_Normal_IGF))
Translation_Normal_fusedData.reset_index(drop=True, inplace=True)

Translation_Normal_fusedmetaData = pd.concat((Translation_Normal_VEH_metaData,Translation_Normal_IGF_metaData))
Translation_Normal_fusedmetaData.reset_index(drop=True, inplace=True)
                                    
Translation_PPHN_fusedData = pd.concat((Translation_PPHN_VEH,Translation_PPHN_IGF))
Translation_PPHN_fusedData.reset_index(drop=True, inplace=True)

Translation_PPHN_fusedmetaData = pd.concat((Translation_PPHN_VEH_metaData,Translation_PPHN_IGF_metaData))
Translation_PPHN_fusedmetaData.reset_index(drop=True, inplace=True)

# %% [code]
# Attempt to normalize data across imaging runs following Carpenter group procedure [pmid:18216396]

#-------------------------------------------------------------------------------
# Find median values, median absolute deviation, and normalize 
# each column in vegf/enos IF data
#-------------------------------------------------------------------------------

# calculate median
IF_Normal_cols = IF_Normal_fusedData.columns
IF_Normal_fusedMedian = IF_Normal_fusedData.median(axis=0).to_frame().T

# Create blank dataframe with feature columns
IF_Normal_fusedNorm = pd.DataFrame(columns=IF_Normal_cols)

# Convert median and median absolute deviaion to rows
IF_Normal_median_data = IF_Normal_fusedMedian.iloc[[0]].values[0]
IF_Normal_mad_data = IF_Normal_dfMAD.iloc[[0]].values[0]

# Subtract median and divide by MAD*1.4826 [pmid:18216396]
IF_Normal_fusedNorm = IF_Normal_fusedData.apply(lambda row: row - IF_Normal_median_data, axis=1, result_type='expand')
IF_Normal_fusedNorm = IF_Normal_fusedNorm.apply(lambda row: row / (IF_Normal_mad_data*1.4826), axis=1, result_type='expand')

# Calculate median
IF_PPHN_cols = IF_PPHN_fusedData.columns
IF_PPHN_fusedMedian = IF_PPHN_fusedData.median(axis=0).to_frame().T

# Create blank dataframe with feature columns
IF_PPHN_fusedNorm = pd.DataFrame(columns=IF_PPHN_cols)

# Convert median and median absolute deviaion to rows
IF_PPHN_median_data = IF_PPHN_fusedMedian.iloc[[0]].values[0]
IF_PPHN_mad_data = IF_PPHN_dfMAD.iloc[[0]].values[0]

# Subtract median and divide by MAD*1.4826 [pmid:18216396]
IF_PPHN_fusedNorm = IF_PPHN_fusedData.apply(lambda row: row - IF_PPHN_median_data, axis=1, result_type='expand')
IF_PPHN_fusedNorm = IF_PPHN_fusedNorm.apply(lambda row: row / (IF_PPHN_mad_data*1.4826), axis=1, result_type='expand')

#-------------------------------------------------------------------------------
# Find median values, median absolute deviation, and normalize 
# each column in translation data
#-------------------------------------------------------------------------------

# calculate median
Translation_Normal_cols = Translation_Normal_fusedData.columns
Translation_Normal_fusedMedian = Translation_Normal_fusedData.median(axis=0).to_frame().T

# Create blank dataframe with feature columns
Translation_Normal_fusedNorm = pd.DataFrame(columns=Translation_Normal_cols)

# Convert median and median absolute deviaion to rows
Translation_Normal_median_data = Translation_Normal_fusedMedian.iloc[[0]].values[0]
Translation_Normal_mad_data = Translation_Normal_dfMAD.iloc[[0]].values[0]

# Subtract median and divide by MAD*1.4826 [pmid:18216396]
Translation_Normal_fusedNorm = Translation_Normal_fusedData.apply(lambda row: row - Translation_Normal_median_data, axis=1, result_type='expand')
Translation_Normal_fusedNorm = Translation_Normal_fusedNorm.apply(lambda row: row / (Translation_Normal_mad_data*1.4826), axis=1, result_type='expand')

# Calculate median
Translation_PPHN_cols = Translation_PPHN_fusedData.columns
Translation_PPHN_fusedMedian = Translation_PPHN_fusedData.median(axis=0).to_frame().T

# Create blank dataframe with feature columns
Translation_PPHN_fusedNorm = pd.DataFrame(columns=Translation_PPHN_cols)

# Convert median and median absolute deviaion to rows
Translation_PPHN_median_data = Translation_PPHN_fusedMedian.iloc[[0]].values[0]
Translation_PPHN_mad_data = Translation_PPHN_dfMAD.iloc[[0]].values[0]

# Subtract median and divide by MAD*1.4826 [pmid:18216396]
Translation_PPHN_fusedNorm = Translation_PPHN_fusedData.apply(lambda row: row - Translation_PPHN_median_data, axis=1, result_type='expand')
Translation_PPHN_fusedNorm = Translation_PPHN_fusedNorm.apply(lambda row: row / (Translation_PPHN_mad_data*1.4826), axis=1, result_type='expand')

# %% [code]
# Transform normalized data across all measurements within each imaging run

#-------------------------------------------------------------------------------
# Calculate z-score of normalized vegf/enos IF data
#-------------------------------------------------------------------------------

IF_Normal_zvalues = StandardScaler().fit_transform(IF_Normal_fusedNorm.values)
IF_Normal_zscored = pd.DataFrame(data = IF_Normal_zvalues, columns=IF_Normal_cols)
IF_Normal_zscored = pd.concat([IF_Normal_zscored, IF_Normal_fusedmetaData], axis = 1)

IF_PPHN_zvalues = StandardScaler().fit_transform(IF_PPHN_fusedNorm.values)
IF_PPHN_zscored = pd.DataFrame(data = IF_PPHN_zvalues, columns=IF_PPHN_cols)
IF_PPHN_zscored = pd.concat([IF_PPHN_zscored, IF_PPHN_fusedmetaData], axis = 1)

#-------------------------------------------------------------------------------
# Calculate z-score for translation data
#-------------------------------------------------------------------------------

Translation_Normal_zvalues = StandardScaler().fit_transform(Translation_Normal_fusedNorm.values)
Translation_Normal_zscored = pd.DataFrame(data = Translation_Normal_zvalues, columns=Translation_Normal_cols)
Translation_Normal_zscored = pd.concat([Translation_Normal_zscored, Translation_Normal_fusedmetaData], axis = 1)

Translation_PPHN_zvalues = StandardScaler().fit_transform(Translation_PPHN_fusedNorm.values)
Translation_PPHN_zscored = pd.DataFrame(data = Translation_PPHN_zvalues, columns=Translation_PPHN_cols)
Translation_PPHN_zscored = pd.concat([Translation_PPHN_zscored, Translation_PPHN_fusedmetaData], axis = 1)


# %% [code]
#-------------------------------------------------------------------------------
# Plot marginal and joint VEGF-eNOS distributions
# This generates the panels for Figures 4 and 5
#-------------------------------------------------------------------------------

sns.set_context("paper", font_scale=2)

times=[0,15,30,60,120,240,1440]

for time in times:
    g=sns.pairplot(IF_Normal_zscored.loc[IF_Normal_zscored['Time']==time], 
                   vars=["Intensity_MeanIntensity_VEGF", "Intensity_MeanIntensity_ENOS"], hue="Media",
                   diag_kind="kde", plot_kws=dict(s=100, alpha=0.3), diag_kws=dict(linewidth=4))
    g.fig.suptitle("Normal PAEC, Time:"+str(time), y=1.08)
    g.axes[0,0].set_xlim((-5,5))
    g.axes[0,0].set_ylabel('VEGF/area')
    g.axes[0,1].set_xlim((-5,5))
    g.axes[1,0].set_xlim((-5,5))
    g.axes[1,0].set_xlabel('eNOS/area')
    g.axes[1,0].set_ylabel('VEGF/area')
    g.axes[1,1].set_xlim((-5,5))
    g.axes[1,1].set_xlabel('eNOS/area')
    g.savefig('figure_04/IF_Normal_PAEC_Time_'+str(time)+'.png', dpi=400)
    
for time in times:
    g=sns.pairplot(IF_PPHN_zscored.loc[IF_PPHN_zscored['Time']==time], 
                   vars=["Intensity_MeanIntensity_VEGF", "Intensity_MeanIntensity_ENOS"], hue="Media", 
                   diag_kind="kde", plot_kws=dict(s=100, alpha=0.3), diag_kws=dict(linewidth=4))
    g.fig.suptitle("PPHN PAEC, Time:"+str(time), y=1.08)
    g.axes[0,0].set_xlim((-5,5))
    g.axes[0,0].set_ylabel('VEGF/area')
    g.axes[0,1].set_xlim((-5,5))
    g.axes[1,0].set_xlim((-5,5))
    g.axes[1,0].set_xlabel('eNOS/area')
    g.axes[1,0].set_ylabel('VEGF/area')
    g.axes[1,1].set_xlim((-5,5))
    g.axes[1,1].set_xlabel('eNOS/area')
    g.savefig('figure_05/IF_PPHN_PAEC_Time_'+str(time)+'.png', dpi=400)

# %% [code]
#-------------------------------------------------------------------------------
# Plot marginal Translation distributions
# This generates Figure 6
#-------------------------------------------------------------------------------

sns.set_context("paper", font_scale=1.5)

g = sns.FacetGrid(Translation_Normal_zscored, col="Time",  row="Cell", hue="Media")
g = (g.map(sns.kdeplot, "Intensity_MeanIntensity_Translation", linewidth=4, shade=True)).add_legend()
g.axes[0,0].set_xlim((-5,5))
g.axes[0,0].set_xlabel('New Protein/area')
g.axes[0,1].set_xlim((-5,5))
g.axes[0,1].set_xlabel('New Protein/area')
g.axes[0,2].set_xlim((-5,5))
g.axes[0,2].set_xlabel('New Protein/area')
g.savefig('figure_06/Translation_Normal_PAEC.png', dpi=400)

g1 = sns.FacetGrid(Translation_PPHN_zscored, col="Time",  row="Cell", hue="Media")
g1 = (g1.map(sns.kdeplot, "Intensity_MeanIntensity_Translation", linewidth=4, shade=True)).add_legend()
g1.axes[0,0].set_xlim((-5,5))
g1.axes[0,0].set_xlabel('New Protein/area')
g1.axes[0,1].set_xlim((-5,5))
g1.axes[0,1].set_xlabel('New Protein/area')
g1.axes[0,2].set_xlim((-5,5))
g1.axes[0,2].set_xlabel('New Protein/area')
g1.savefig('figure_06/Translation_PPHN_PAEC.png', dpi=400)

# %% [code]
# Perform PCA on z-scored data

# Set number of principal components to capture 95% of the variance
numberPC=115

# Calculate PCA
IF_Normal_pca=PCA(n_components=numberPC)
IF_Normal_principalComponents=IF_Normal_pca.fit_transform(IF_Normal_zvalues)
IF_Normal_principal = pd.DataFrame(data = IF_Normal_principalComponents)

# Uncomment this to output how much variance is captured by given number of principal components
#print(np.sum(pca.explained_variance_ratio_))

# Generate column names for PCA
pcColNames=[]
for i in range(1,numberPC+1):
    pcColNames.append("PC"+ str(i))

# Merge PCA per cell with metadata
IF_Normal_PCA = pd.concat([IF_Normal_principal, IF_Normal_fusedmetaData], axis = 1)

# Calculate PCA
IF_PPHN_pca=PCA(n_components=numberPC)
IF_PPHN_principalComponents=IF_PPHN_pca.fit_transform(IF_PPHN_zvalues)
IF_PPHN_principal = pd.DataFrame(data = IF_PPHN_principalComponents)

# Uncomment this to output how much variance is captured by given number of principal components
#print(np.sum(pca.explained_variance_ratio_))

# Generate column names for PCA
pcColNames=[]
for i in range(1,numberPC+1):
    pcColNames.append("PC"+ str(i))

# Merge PCA per cell with metadata
IF_PPHN_PCA = pd.concat([IF_PPHN_principal, IF_PPHN_fusedmetaData], axis = 1)

# Calculate PCA
Translation_Normal_pca=PCA(n_components=numberPC)
Translation_Normal_principalComponents=Translation_Normal_pca.fit_transform(Translation_Normal_zvalues)
Translation_Normal_principal = pd.DataFrame(data = Translation_Normal_principalComponents)

# Uncomment this to output how much variance is captured by given number of principal components
#print(np.sum(pca.explained_variance_ratio_))

# Generate column names for PCA
pcColNames=[]
for i in range(1,numberPC+1):
    pcColNames.append("PC"+ str(i))

# Merge PCA per cell with metadata
Translation_Normal_PCA = pd.concat([Translation_Normal_principal, Translation_Normal_fusedmetaData], axis = 1)

# Calculate PCA
Translation_PPHN_pca=PCA(n_components=numberPC)
Translation_PPHN_principalComponents=Translation_PPHN_pca.fit_transform(Translation_PPHN_zvalues)
Translation_PPHN_principal = pd.DataFrame(data = Translation_PPHN_principalComponents)

# Uncomment this to output how much variance is captured by given number of principal components
#print(np.sum(pca.explained_variance_ratio_))

# Generate column names for PCA
pcColNames=[]
for i in range(1,numberPC+1):
    pcColNames.append("PC"+ str(i))

# Merge PCA per cell with metadata
Translation_PPHN_PCA = pd.concat([Translation_PPHN_principal, Translation_PPHN_fusedmetaData], axis = 1)

# %% [code]
# Virtually stitch images for use in spatial analysis and assign an identifier to each replicate
# Images were acquired with 0% overlap, so we can just add the camera chip size (2048 x 2048 pixels) in the correct
# order based on the MicroManager tiling order.


#-------------------------------------------------------------------------------
# Tile images with PCA data for normal PAEC vegf/enos
#-------------------------------------------------------------------------------
# Define conditions
cells=['Normal']
times=[0, 15, 30, 60, 120, 240, 1440]
medias=['Vehicle', 'IGF1']

# Create blank dataframe to hold tiled data
IF_Normal_tiled_PCA=pd.DataFrame(columns=IF_Normal_PCA.columns)

for time in times:
    for cell in cells:
        for media in medias:
            # Pull data for current condition
            dfTemp=IF_Normal_PCA[(IF_Normal_PCA['Cell']==cell) & (IF_Normal_PCA['Time']==time) & (IF_Normal_PCA['Media']==media)]
            dfTemp.reset_index(drop=True, inplace=True)
            
            # Reset image numbers to 1 -> 100 within a given condition
            if media == 'Vehicle':
                Vehicle_index =dfTemp[dfTemp.loc[:,'Media']=='Vehicle'].index
                dfTemp.loc[Vehicle_index,'ImageNumber'] = dfTemp.loc[Vehicle_index, 'ImageNumber'].sub(dfTemp.loc[Vehicle_index,'ImageNumber'].min())+1
            else:
                IGF1_index = dfTemp[dfTemp.loc[:, "Media"] == "IGF1"].index
                dfTemp.loc[IGF1_index,'ImageNumber'] = dfTemp.loc[IGF1_index,'ImageNumber'].sub(dfTemp.loc[IGF1_index,'ImageNumber'].min())+1
                dfTemp.loc[IGF1_index, "AreaShape_Center_Y"] = dfTemp.loc[IGF1_index, "AreaShape_Center_Y"]+11*2100
            
            # Create dummy dataframe for storage
            dfTemp3=pd.DataFrame(columns=dfTemp.columns)
            dfTemp3.drop(['ImageNumber'],axis=1,inplace=True)
                        
            # Counter for row location in tiled area
            counter = 0
            
            # Stitch images based on MicroManager tile order 
            for image in range(1,200):
                
                # Find all cells in the current image
                idx = dfTemp.index[dfTemp['ImageNumber']==image]
            
                # Pull data from current image and drop Image Number
                dfTemp2=dfTemp.loc[idx]
                dfTemp2.drop(['ImageNumber'],axis=1,inplace=True)
            
                # Move cells in (x,y) according to tile and image position
                if image <= 100:
                    dfTemp2['AreaShape_Center_X']=dfTemp['AreaShape_Center_X']+(counter)*2048
                    dfTemp2['AreaShape_Center_Y']=dfTemp['AreaShape_Center_Y']+(np.floor_divide(image-1,10)*2048)
                    dfTemp2['Replicate']=1
                else:
                    dfTemp2['AreaShape_Center_X']=dfTemp['AreaShape_Center_X']+(counter)*2048+(10*2048)
                    dfTemp2['AreaShape_Center_Y']=dfTemp['AreaShape_Center_Y']+(np.floor_divide(image-1,10)*2048-(10*2048))
                    dfTemp2['Replicate']=2
                counter = counter + 1
                if counter == 9:
                    counter = 0
                    
                # Add data from current image to dummy dataframe for tiled area
                dfTemp3=pd.concat([dfTemp3,dfTemp2],ignore_index=True)
                
            # Reset index of tiled area
            dfTemp3.reset_index(drop=True, inplace=True)
            
            # Reset object counter
            dfTemp3.ObjectNumber = np.arange(1,len(dfTemp3)+1)
        
            # Add results to final dataframe
            IF_Normal_tiled_PCA=pd.concat([IF_Normal_tiled_PCA,dfTemp3],ignore_index=True)

# Clean up final dataframe
IF_Normal_tiled_PCA.drop(['ImageNumber'],axis=1,inplace=True)


#-------------------------------------------------------------------------------
# Tile images with PCA data for PPHN PAEC vegf/enos
#-------------------------------------------------------------------------------

cells=['PPHN']
times=[0, 15, 30, 60, 120, 240, 1440]
medias=['Vehicle', 'IGF1']


IF_PPHN_tiled_PCA=pd.DataFrame(columns=IF_PPHN_PCA.columns)

for time in times:
    for cell in cells:
        for media in medias:
            # Pull data for current condition
            dfTemp=IF_PPHN_PCA[(IF_PPHN_PCA['Cell']==cell) & (IF_PPHN_PCA['Time']==time) & (IF_PPHN_PCA['Media']==media)]
            dfTemp.reset_index(drop=True, inplace=True)
            
            # Reset image numbers to 1 -> 100 within a given condition
            if media == 'Vehicle':
                Vehicle_index =dfTemp[dfTemp.loc[:,'Media']=='Vehicle'].index
                dfTemp.loc[Vehicle_index,'ImageNumber'] = dfTemp.loc[Vehicle_index, 'ImageNumber'].sub(dfTemp.loc[Vehicle_index,'ImageNumber'].min())+1
            else:
                IGF1_index = dfTemp[dfTemp.loc[:, "Media"] == "IGF1"].index
                dfTemp.loc[IGF1_index,'ImageNumber'] = dfTemp.loc[IGF1_index,'ImageNumber'].sub(dfTemp.loc[IGF1_index,'ImageNumber'].min())+1
                dfTemp.loc[IGF1_index, "AreaShape_Center_Y"] = dfTemp.loc[IGF1_index, "AreaShape_Center_Y"]+11*2100
            
            # Create dummy dataframe for storage
            dfTemp3=pd.DataFrame(columns=dfTemp.columns)
            dfTemp3.drop(['ImageNumber'],axis=1,inplace=True)
                        
            # Counter for row location in tiled area
            counter = 0
            
            # Stitch images based on MicroManager tile order 
            for image in range(1,200):
                
                # Find all cells in the current image
                idx = dfTemp.index[dfTemp['ImageNumber']==image]
            
                # Pull data from current image and drop Image Number
                dfTemp2=dfTemp.loc[idx]
                dfTemp2.drop(['ImageNumber'],axis=1,inplace=True)
            
                # Move cells in (x,y) according to tile and image position
                if image <= 100:
                    dfTemp2['AreaShape_Center_X']=dfTemp['AreaShape_Center_X']+(counter)*2048
                    dfTemp2['AreaShape_Center_Y']=dfTemp['AreaShape_Center_Y']+(np.floor_divide(image-1,10)*2048)
                    dfTemp2['Replicate']=1
                else:
                    dfTemp2['AreaShape_Center_X']=dfTemp['AreaShape_Center_X']+(counter)*2048+(10*2048)
                    dfTemp2['AreaShape_Center_Y']=dfTemp['AreaShape_Center_Y']+(np.floor_divide(image-1,10)*2048-(10*2048))
                    dfTemp2['Replicate']=2
                counter = counter + 1
                if counter == 9:
                    counter = 0
                    
                # Add data from current image to dummy dataframe for tiled area
                dfTemp3=pd.concat([dfTemp3,dfTemp2],ignore_index=True)
                
            # Reset index of tiled area
            dfTemp3.reset_index(drop=True, inplace=True)
            
            # Reset object counter
            dfTemp3.ObjectNumber = np.arange(1,len(dfTemp3)+1)
        
            # Add results to final dataframe
            IF_PPHN_tiled_PCA=pd.concat([IF_PPHN_tiled_PCA,dfTemp3],ignore_index=True)

# Clean up final dataframe
IF_PPHN_tiled_PCA.drop(['ImageNumber'],axis=1,inplace=True)

#-------------------------------------------------------------------------------
# Tile images with PCA data for Normal PAEC translation
#-------------------------------------------------------------------------------

cells=['Normal']
times=[0, 60, 1440]
medias=['Vehicle', 'IGF1']

Translation_Normal_tiled_PCA=pd.DataFrame(columns=Translation_Normal_PCA.columns)

for time in times:
    for cell in cells:
        for media in medias:
            # Pull data for current condition
            dfTemp=Translation_Normal_PCA[(Translation_Normal_PCA['Cell']==cell) & (Translation_Normal_PCA['Time']==time) & (Translation_Normal_PCA['Media']==media)]
            dfTemp.reset_index(drop=True, inplace=True)
            
            # Reset image numbers to 1 -> 100 within a given condition
            if media == 'Vehicle':
                Vehicle_index =dfTemp[dfTemp.loc[:,'Media']=='Vehicle'].index
                dfTemp.loc[Vehicle_index,'ImageNumber'] = dfTemp.loc[Vehicle_index, 'ImageNumber'].sub(dfTemp.loc[Vehicle_index,'ImageNumber'].min())+1
            else:
                IGF1_index = dfTemp[dfTemp.loc[:, "Media"] == "IGF1"].index
                dfTemp.loc[IGF1_index,'ImageNumber'] = dfTemp.loc[IGF1_index,'ImageNumber'].sub(dfTemp.loc[IGF1_index,'ImageNumber'].min())+1
                dfTemp.loc[IGF1_index, "AreaShape_Center_Y"] = dfTemp.loc[IGF1_index, "AreaShape_Center_Y"]+11*2100
            
            # Create dummy dataframe for storage
            dfTemp3=pd.DataFrame(columns=dfTemp.columns)
            dfTemp3.drop(['ImageNumber'],axis=1,inplace=True)
                        
            # Counter for row location in tiled area
            counter = 0
            
            # Stitch images based on MicroManager tile order 
            for image in range(1,200):
                
                # Find all cells in the current image
                idx = dfTemp.index[dfTemp['ImageNumber']==image]
            
                # Pull data from current image and drop Image Number
                dfTemp2=dfTemp.loc[idx]
                dfTemp2.drop(['ImageNumber'],axis=1,inplace=True)
            
                # Move cells in (x,y) according to tile and image position
                if image <= 100:
                    dfTemp2['AreaShape_Center_X']=dfTemp['AreaShape_Center_X']+(counter)*2048
                    dfTemp2['AreaShape_Center_Y']=dfTemp['AreaShape_Center_Y']+(np.floor_divide(image-1,10)*2048)
                    dfTemp2['Replicate']=1
                else:
                    dfTemp2['AreaShape_Center_X']=dfTemp['AreaShape_Center_X']+(counter)*2048+(10*2048)
                    dfTemp2['AreaShape_Center_Y']=dfTemp['AreaShape_Center_Y']+(np.floor_divide(image-1,10)*2048-(10*2048))
                    dfTemp2['Replicate']=2
                counter = counter + 1
                if counter == 9:
                    counter = 0
                    
                # Add data from current image to dummy dataframe for tiled area
                dfTemp3=pd.concat([dfTemp3,dfTemp2],ignore_index=True)
                
            # Reset index of tiled area
            dfTemp3.reset_index(drop=True, inplace=True)
            
            # Reset object counter
            dfTemp3.ObjectNumber = np.arange(1,len(dfTemp3)+1)
        
            # Add results to final dataframe
            Translation_Normal_tiled_PCA=pd.concat([Translation_Normal_tiled_PCA,dfTemp3],ignore_index=True)

# Clean up final dataframe
Translation_Normal_tiled_PCA.drop(['ImageNumber'],axis=1,inplace=True)

#-------------------------------------------------------------------------------
# Tile images with PCA data for PPHN PAEC translation
#-------------------------------------------------------------------------------
cells=['PPHN']
times=[0, 60, 1440]
medias=['Vehicle', 'IGF1']

Translation_PPHN_tiled_PCA=pd.DataFrame(columns=Translation_PPHN_PCA.columns)

for time in times:
    for cell in cells:
        for media in medias:
            # Pull data for current condition
            dfTemp=Translation_PPHN_PCA[(Translation_PPHN_PCA['Cell']==cell) & (Translation_PPHN_PCA['Time']==time) & (Translation_PPHN_PCA['Media']==media)]
            dfTemp.reset_index(drop=True, inplace=True)
            
            # Reset image numbers to 1 -> 100 within a given condition
            if media == 'Vehicle':
                Vehicle_index =dfTemp[dfTemp.loc[:,'Media']=='Vehicle'].index
                dfTemp.loc[Vehicle_index,'ImageNumber'] = dfTemp.loc[Vehicle_index, 'ImageNumber'].sub(dfTemp.loc[Vehicle_index,'ImageNumber'].min())+1
            else:
                IGF1_index = dfTemp[dfTemp.loc[:, "Media"] == "IGF1"].index
                dfTemp.loc[IGF1_index,'ImageNumber'] = dfTemp.loc[IGF1_index,'ImageNumber'].sub(dfTemp.loc[IGF1_index,'ImageNumber'].min())+1
                dfTemp.loc[IGF1_index, "AreaShape_Center_Y"] = dfTemp.loc[IGF1_index, "AreaShape_Center_Y"]+11*2100
            
            # Create dummy dataframe for storage
            dfTemp3=pd.DataFrame(columns=dfTemp.columns)
            dfTemp3.drop(['ImageNumber'],axis=1,inplace=True)
                        
            # Counter for row location in tiled area
            counter = 0
            
            # Stitch images based on MicroManager tile order 
            for image in range(1,200):
                
                # Find all cells in the current image
                idx = dfTemp.index[dfTemp['ImageNumber']==image]
            
                # Pull data from current image and drop Image Number
                dfTemp2=dfTemp.loc[idx]
                dfTemp2.drop(['ImageNumber'],axis=1,inplace=True)
            
                # Move cells in (x,y) according to tile and image position
                if image <= 100:
                    dfTemp2['AreaShape_Center_X']=dfTemp['AreaShape_Center_X']+(counter)*2048
                    dfTemp2['AreaShape_Center_Y']=dfTemp['AreaShape_Center_Y']+(np.floor_divide(image-1,10)*2048)
                    dfTemp2['Replicate']=1
                else:
                    dfTemp2['AreaShape_Center_X']=dfTemp['AreaShape_Center_X']+(counter)*2048+(10*2048)
                    dfTemp2['AreaShape_Center_Y']=dfTemp['AreaShape_Center_Y']+(np.floor_divide(image-1,10)*2048-(10*2048))
                    dfTemp2['Replicate']=2
                counter = counter + 1
                if counter == 9:
                    counter = 0
                    
                # Add data from current image to dummy dataframe for tiled area
                dfTemp3=pd.concat([dfTemp3,dfTemp2],ignore_index=True)
                
            # Reset index of tiled area
            dfTemp3.reset_index(drop=True, inplace=True)
            
            # Reset object counter
            dfTemp3.ObjectNumber = np.arange(1,len(dfTemp3)+1)
        
            # Add results to final dataframe
            Translation_PPHN_tiled_PCA=pd.concat([Translation_PPHN_tiled_PCA,dfTemp3],ignore_index=True)

# Clean up final dataframe
Translation_PPHN_tiled_PCA.drop(['ImageNumber'],axis=1,inplace=True)

# %% [code]
# Calculate the null distribution of Pearson correlations within one condition/replicates by ignoring spatial coordinates
# We randomly pull 2000 pairs of cells in a given condition and calculate the Pearson correlation between them

#-------------------------------------------------------------------------------
# Calculate null distribution for normal PAEC vegf/enos
#-------------------------------------------------------------------------------

# Define conditions
cells=['Normal']
times=[0,15,30,60,120,240,1440]
medias=['Vehicle','IGF1']
replicates=[1,2]

IF_Normal_NullCellCorr=pd.DataFrame(columns=['Cell type','Media type','Time point','Correlation','Replicate'])

for time in times:
    for cell in cells:
        for media in medias:
            for replicate in replicates:

                nullCellCorr=np.zeros(1999)

                # Create blank dataframe for storage of all replicates
                dfNullCellCorrCalc=pd.DataFrame(columns=IF_Normal_tiled_PCA.columns)
                dfNullCellCorrCalc=dfNullCellCorrCalc.fillna(0)

                # Find maximum number of objects for condition
                dfTemp=IF_Normal_tiled_PCA[(IF_Normal_tiled_PCA["Cell"]==cell) & (IF_Normal_tiled_PCA["Media"]==media) 
                                 & (IF_Normal_tiled_PCA["Time"]==time)
                                 & (IF_Normal_tiled_PCA["Replicate"]==replicate)]
                dfTemp.dropna(axis=0,inplace=True)
                dfTemp.reset_index(inplace=True)
                dfTemp.drop(['index'],axis=1,inplace=True)
                maxObjects=dfTemp.shape[0]

                # Calculate correlation for 2000 random cell pairs
                for i in range(0,1999):

                    # Pull two random cells from replicate
                    randObject_1=random.randint(0,maxObjects-1)
                    randObject_2=random.randint(0,maxObjects-1)

                    # Assign featres from cells to dataframe
                    dfTemp2=dfTemp.iloc[[randObject_1]]
                    dfTemp3=dfTemp.iloc[[randObject_2]]
                    dfNullCellCorrCalc=pd.concat([dfTemp2,dfTemp3],ignore_index=True)

                    # calculate correlation between cells features 
                    dfNullCellCorrCalc.drop(['Cell','Media','Time','ObjectNumber','Replicate',
                                             'AreaShape_Center_X','AreaShape_Center_Y'],axis=1,inplace=True)
                    dfNullCellCorrCalc=dfNullCellCorrCalc.T
                    dfNullCellCorrMatrix=dfNullCellCorrCalc.corr()
                    nullCellCorr[i]=dfNullCellCorrMatrix[0][1]

                # Add all pair correlations to final dataframe
                dfTempNullCorr=pd.DataFrame(columns=['Cell type','Media type','Time point','Correlation','Replicate'])
                dfTempNullCorr['Correlation']=nullCellCorr
                dfTempNullCorr['Cell type']=cell
                dfTempNullCorr['Media type']=media
                dfTempNullCorr['Time point']=time
                dfTempNullCorr['Replicate']=replicate

                IF_Normal_NullCellCorr=pd.concat([IF_Normal_NullCellCorr,dfTempNullCorr],ignore_index=True)
                
#-------------------------------------------------------------------------------
# Calculate null distribution for PPHN PAEC vegf/enos
#-------------------------------------------------------------------------------

# Define conditions
cells=['PPHN']
times=[0,15,30,60,120,240,1440]
medias=['Vehicle','IGF1']
replicates=[1,2]

IF_PPHN_NullCellCorr=pd.DataFrame(columns=['Cell type','Media type','Time point','Correlation','Replicate'])

for time in times:
    for cell in cells:
        for media in medias:
            for replicate in replicates:

                nullCellCorr=np.zeros(1999)

                # Create blank dataframe for storage of all replicates
                dfNullCellCorrCalc=pd.DataFrame(columns=IF_PPHN_tiled_PCA.columns)
                dfNullCellCorrCalc=dfNullCellCorrCalc.fillna(0)

                # Fix missing replicates for PPHN data.
                # This won't be used in downstream analysis, it's just easier to do this way.
                if ((media=='Vehicle') & (time==120) & (replicate==2)):
                    replicate=1
                    
                if ((media=='IGF1') & (time==240) & (replicate==2)):
                    replicate=1
                    
                # Find maximum number of objects for condition
                dfTemp=IF_PPHN_tiled_PCA[(IF_PPHN_tiled_PCA["Cell"]==cell) & (IF_PPHN_tiled_PCA["Media"]==media) 
                                 & (IF_PPHN_tiled_PCA["Time"]==time)
                                 & (IF_PPHN_tiled_PCA["Replicate"]==replicate)]
                dfTemp.dropna(axis=0,inplace=True)
                dfTemp.reset_index(inplace=True)
                dfTemp.drop(['index'],axis=1,inplace=True)
                maxObjects=dfTemp.shape[0]
         
                # Calculate correlation for 2000 random cell pairs
                for i in range(0,1999):

                    # Pull two random cells from replicate
                    randObject_1=random.randint(0,maxObjects-1)
                    randObject_2=random.randint(0,maxObjects-1)

                    # Assign featres from cells to dataframe
                    dfTemp2=dfTemp.iloc[[randObject_1]]
                    dfTemp3=dfTemp.iloc[[randObject_2]]
                    dfNullCellCorrCalc=pd.concat([dfTemp2,dfTemp3],ignore_index=True)

                    # calculate correlation between cells features 
                    dfNullCellCorrCalc.drop(['Cell','Media','Time','ObjectNumber','Replicate',
                                             'AreaShape_Center_X','AreaShape_Center_Y'],axis=1,inplace=True)
                    dfNullCellCorrCalc=dfNullCellCorrCalc.T
                    dfNullCellCorrMatrix=dfNullCellCorrCalc.corr()
                    nullCellCorr[i]=dfNullCellCorrMatrix[0][1]

                # Add all pair correlations to final dataframe
                dfTempNullCorr=pd.DataFrame(columns=['Cell type','Media type','Time point','Correlation','Replicate'])
                dfTempNullCorr['Correlation']=nullCellCorr
                dfTempNullCorr['Cell type']=cell
                dfTempNullCorr['Media type']=media
                dfTempNullCorr['Time point']=time
                dfTempNullCorr['Replicate']=replicate

                IF_PPHN_NullCellCorr=pd.concat([IF_PPHN_NullCellCorr,dfTempNullCorr],ignore_index=True)
                
                
#-------------------------------------------------------------------------------
# Calculate null distribution for Normal PAEC translation
#-------------------------------------------------------------------------------

# Define conditions
cells=['Normal']
times=[0,60,1440]
medias=['Vehicle','IGF1']
replicates=[1,2]

Translation_Normal_NullCellCorr=pd.DataFrame(columns=['Cell type','Media type','Time point','Correlation','Replicate'])

for time in times:
    for cell in cells:
        for media in medias:
            for replicate in replicates:

                nullCellCorr=np.zeros(1999)

                # Create blank dataframe for storage of all replicates
                dfNullCellCorrCalc=pd.DataFrame(columns=Translation_Normal_tiled_PCA.columns)
                dfNullCellCorrCalc=dfNullCellCorrCalc.fillna(0)

                # Find maximum number of objects for condition
                dfTemp=Translation_Normal_tiled_PCA[(Translation_Normal_tiled_PCA["Cell"]==cell) 
                                                   & (Translation_Normal_tiled_PCA["Media"]==media) 
                                                   & (Translation_Normal_tiled_PCA["Time"]==time)
                                                   & (Translation_Normal_tiled_PCA["Replicate"]==replicate)]
                dfTemp.dropna(axis=0,inplace=True)
                dfTemp.reset_index(inplace=True)
                dfTemp.drop(['index'],axis=1,inplace=True)
                maxObjects=dfTemp.shape[0]

                # Calculate correlation for 2000 random cell pairs
                for i in range(0,1999):

                    # Pull two random cells from replicate
                    randObject_1=random.randint(0,maxObjects-1)
                    randObject_2=random.randint(0,maxObjects-1)

                    # Assign featres from cells to dataframe
                    dfTemp2=dfTemp.iloc[[randObject_1]]
                    dfTemp3=dfTemp.iloc[[randObject_2]]
                    dfNullCellCorrCalc=pd.concat([dfTemp2,dfTemp3],ignore_index=True)

                    # calculate correlation between cells features 
                    dfNullCellCorrCalc.drop(['Cell','Media','Time','ObjectNumber','Replicate',
                                             'AreaShape_Center_X','AreaShape_Center_Y'],axis=1,inplace=True)
                    dfNullCellCorrCalc=dfNullCellCorrCalc.T
                    dfNullCellCorrMatrix=dfNullCellCorrCalc.corr()
                    nullCellCorr[i]=dfNullCellCorrMatrix[0][1]

                # Add all pair correlations to final dataframe
                dfTempNullCorr=pd.DataFrame(columns=['Cell type','Media type','Time point','Correlation','Replicate'])
                dfTempNullCorr['Correlation']=nullCellCorr
                dfTempNullCorr['Cell type']=cell
                dfTempNullCorr['Media type']=media
                dfTempNullCorr['Time point']=time
                dfTempNullCorr['Replicate']=replicate

                Translation_Normal_NullCellCorr=pd.concat([Translation_Normal_NullCellCorr,dfTempNullCorr],ignore_index=True)
                
#-------------------------------------------------------------------------------
# Calculate null distribution for PPHN PAEC translation
#-------------------------------------------------------------------------------

# Define conditions
cells=['PPHN']
times=[0,60,1440]
medias=['Vehicle','IGF1']
replicates=[1,2]

Translation_PPHN_NullCellCorr=pd.DataFrame(columns=['Cell type','Media type','Time point','Correlation','Replicate'])

for time in times:
    for cell in cells:
        for media in medias:
            for replicate in replicates:

                nullCellCorr=np.zeros(1999)

                # Create blank dataframe for storage of all replicates
                dfNullCellCorrCalc=pd.DataFrame(columns=Translation_PPHN_tiled_PCA.columns)
                dfNullCellCorrCalc=dfNullCellCorrCalc.fillna(0)

                # Find maximum number of objects for condition
                dfTemp=Translation_PPHN_tiled_PCA[(Translation_PPHN_tiled_PCA["Cell"]==cell) 
                                                   & (Translation_PPHN_tiled_PCA["Media"]==media) 
                                                   & (Translation_PPHN_tiled_PCA["Time"]==time)
                                                   & (Translation_PPHN_tiled_PCA["Replicate"]==replicate)]
                dfTemp.dropna(axis=0,inplace=True)
                dfTemp.reset_index(inplace=True)
                dfTemp.drop(['index'],axis=1,inplace=True)
                maxObjects=dfTemp.shape[0]

                # Calculate correlation for 2000 random cell pairs
                for i in range(0,1999):

                    # Pull two random cells from replicate
                    randObject_1=random.randint(0,maxObjects-1)
                    randObject_2=random.randint(0,maxObjects-1)

                    # Assign featres from cells to dataframe
                    dfTemp2=dfTemp.iloc[[randObject_1]]
                    dfTemp3=dfTemp.iloc[[randObject_2]]
                    dfNullCellCorrCalc=pd.concat([dfTemp2,dfTemp3],ignore_index=True)

                    # calculate correlation between cells features 
                    dfNullCellCorrCalc.drop(['Cell','Media','Time','ObjectNumber','Replicate',
                                             'AreaShape_Center_X','AreaShape_Center_Y'],axis=1,inplace=True)
                    dfNullCellCorrCalc=dfNullCellCorrCalc.T
                    dfNullCellCorrMatrix=dfNullCellCorrCalc.corr()
                    nullCellCorr[i]=dfNullCellCorrMatrix[0][1]

                # Add all pair correlations to final dataframe
                dfTempNullCorr=pd.DataFrame(columns=['Cell type','Media type','Time point','Correlation','Replicate'])
                dfTempNullCorr['Correlation']=nullCellCorr
                dfTempNullCorr['Cell type']=cell
                dfTempNullCorr['Media type']=media
                dfTempNullCorr['Time point']=time
                dfTempNullCorr['Replicate']=replicate

                Translation_PPHN_NullCellCorr=pd.concat([Translation_Normal_NullCellCorr,dfTempNullCorr],ignore_index=True)

# %% [code]
# Calculate the spatial Pearson correlations within one condition/replicates
# Pull the 20 nearest neighbors for each cell and calculate the median Pearson correlation coefficient

#-------------------------------------------------------------------------------
# Calculate spatial correlation for normal PAEC vegf/enos
#-------------------------------------------------------------------------------

# Setup conditions
cells=['Normal']
times=[0,15,30,60,120,240,1440]
medias=['Vehicle','IGF1']
replicates=[1,2]

IF_Normal_SpatialCellCorr=pd.DataFrame(columns=['Cell type','Media type','Time point','Correlation', 'xPos','yPos'])

for time in times:
    for cell in cells:
        for media in medias:
            for replicate in replicates:

                dfTemp=IF_Normal_tiled_PCA[(IF_Normal_tiled_PCA["Cell"]==cell) & (IF_Normal_tiled_PCA["Media"]==media) 
                                 & (IF_Normal_tiled_PCA["Time"]==time)
                                 & (IF_Normal_tiled_PCA["Replicate"]==replicate)]
                dfTemp.dropna(axis=0,inplace=True)
                dfTemp.reset_index(inplace=True)
                dfTemp.drop(['index'],axis=1,inplace=True)
                maxObjects=dfTemp.shape[0]
                maxCorrelation=20

                cellCorr=np.zeros((maxObjects,maxCorrelation))
                xPos=np.zeros(maxObjects)
                yPos=np.zeros(maxObjects)

                for i in range(0,maxObjects):
                    dfCurrentCell=dfTemp.iloc[[i]]
                    dfCellCorrCalc=dfCurrentCell
                    distanceToCells=np.sqrt((dfCurrentCell['AreaShape_Center_X'].values-dfTemp['AreaShape_Center_X'].values)**2+
                                                    (dfCurrentCell['AreaShape_Center_Y'].values-dfTemp['AreaShape_Center_Y'].values)**2)
                    tempList=distanceToCells[1:-1]
                    nearestCells=sorted(np.argpartition(tempList, maxCorrelation)[:maxCorrelation])
                    xPos[i]=dfCurrentCell['AreaShape_Center_X']
                    yPos[i]=dfCurrentCell['AreaShape_Center_Y']
                    for j in range(0,maxCorrelation):
                        dfCellCorrCalc=pd.concat([dfCellCorrCalc,dfTemp.iloc[[nearestCells[j]]]],ignore_index=True)
                        dfCellCorrCalc.drop(['Cell','Media','Time','ObjectNumber',
                                        'AreaShape_Center_X','AreaShape_Center_Y'],axis=1,inplace=True)
                        dfCellCorrCalcTranspose=dfCellCorrCalc.T
                        dfCellCorrMatrix=dfCellCorrCalcTranspose.corr()
                        if j > 0:
                            medianCorr=np.round(dfCellCorrMatrix[1:-1].median(axis=0),2)
                            cellCorr[i][j]=medianCorr[0]
                        else:
                            cellCorr[i][j]=np.round(dfCellCorrMatrix[0][1],2)

                # Add all median spatial correlations to final dataframe
                medianCorr=np.round(np.median(cellCorr,axis=1),2)
                dfTempCorr=pd.DataFrame(columns=['Cell type','Media type','Time point','Correlation'])
                dfTempCorr['Correlation']=medianCorr
                dfTempCorr['Cell type']=cell
                dfTempCorr['Media type']=media
                dfTempCorr['Time point']=time
                dfTempCorr['xPos']=xPos
                dfTempCorr['yPos']=yPos
                dfTempCorr['Replicate']=replicate

                IF_Normal_SpatialCellCorr=pd.concat([IF_Normal_SpatialCellCorr,dfTempCorr],ignore_index=True)
                
#-------------------------------------------------------------------------------
#  Calculate spatial correlation for PPHN PAEC vegf/enos
#-------------------------------------------------------------------------------

# Setup conditions
cells=['PPHN']
times=[0,15,30,60,120,240,1440]
medias=['Vehicle','IGF1']
replicates=[1,2]

IF_PPHN_SpatialCellCorr=pd.DataFrame(columns=['Cell type','Media type','Time point','Correlation', 'xPos','yPos'])

for time in times:
    for cell in cells:
        for media in medias:
            for replicate in replicates:
                
                # Fix missing replicates for PPHN data.
                # This won't be used in downstream analysis, it's just easier to do this way.
                if ((media=='Vehicle') & (time==120) & (replicate==2)):
                    replicate=1
                    
                if ((media=='IGF1') & (time==240) & (replicate==2)):
                    replicate=1

                dfTemp=IF_PPHN_tiled_PCA[(IF_PPHN_tiled_PCA["Cell"]==cell) & (IF_PPHN_tiled_PCA["Media"]==media) 
                                 & (IF_PPHN_tiled_PCA["Time"]==time)
                                 & (IF_PPHN_tiled_PCA["Replicate"]==replicate)]
                dfTemp.dropna(axis=0,inplace=True)
                dfTemp.reset_index(inplace=True)
                dfTemp.drop(['index'],axis=1,inplace=True)
                maxObjects=dfTemp.shape[0]
                maxCorrelation=20

                cellCorr=np.zeros((maxObjects,maxCorrelation))
                xPos=np.zeros(maxObjects)
                yPos=np.zeros(maxObjects)

                for i in range(0,maxObjects):
                    dfCurrentCell=dfTemp.iloc[[i]]
                    dfCellCorrCalc=dfCurrentCell
                    distanceToCells=np.sqrt((dfCurrentCell['AreaShape_Center_X'].values-dfTemp['AreaShape_Center_X'].values)**2+
                                                    (dfCurrentCell['AreaShape_Center_Y'].values-dfTemp['AreaShape_Center_Y'].values)**2)
                    tempList=distanceToCells[1:-1]
                    nearestCells=sorted(np.argpartition(tempList, maxCorrelation)[:maxCorrelation])
                    xPos[i]=dfCurrentCell['AreaShape_Center_X']
                    yPos[i]=dfCurrentCell['AreaShape_Center_Y']
                    for j in range(0,maxCorrelation):
                        dfCellCorrCalc=pd.concat([dfCellCorrCalc,dfTemp.iloc[[nearestCells[j]]]],ignore_index=True)
                        dfCellCorrCalc.drop(['Cell','Media','Time','ObjectNumber',
                                        'AreaShape_Center_X','AreaShape_Center_Y'],axis=1,inplace=True)
                        dfCellCorrCalcTranspose=dfCellCorrCalc.T
                        dfCellCorrMatrix=dfCellCorrCalcTranspose.corr()
                        if j > 0:
                            medianCorr=np.round(dfCellCorrMatrix[1:-1].median(axis=0),2)
                            cellCorr[i][j]=medianCorr[0]
                        else:
                            cellCorr[i][j]=np.round(dfCellCorrMatrix[0][1],2)

                # Add all median spatial correlations to final dataframe
                medianCorr=np.round(np.median(cellCorr,axis=1),2)
                dfTempCorr=pd.DataFrame(columns=['Cell type','Media type','Time point','Correlation'])
                dfTempCorr['Correlation']=medianCorr
                dfTempCorr['Cell type']=cell
                dfTempCorr['Media type']=media
                dfTempCorr['Time point']=time
                dfTempCorr['xPos']=xPos
                dfTempCorr['yPos']=yPos
                dfTempCorr['Replicate']=replicate

                IF_PPHN_SpatialCellCorr=pd.concat([IF_PPHN_SpatialCellCorr,dfTempCorr],ignore_index=True)    
                
#-------------------------------------------------------------------------------
#  Calculate spatial correlation for Normal PAEC translation
#-------------------------------------------------------------------------------

# Setup conditions
cells=['Normal']
times=[0,15,30,60,120,240,1440]
medias=['Vehicle','IGF1']
replicates=[1,2]

Translation_Normal_SpatialCellCorr=pd.DataFrame(columns=['Cell type','Media type','Time point','Correlation', 'xPos','yPos'])

for time in times:
    for cell in cells:
        for media in medias:
            for replicate in replicates:
                
                # Fix missing replicates for PPHN data.
                # This won't be used in downstream analysis, it's just easier to do this way.
                if ((media=='Vehicle') & (time==120) & (replicate==2)):
                    replicate=1
                    
                if ((media=='IGF1') & (time==240) & (replicate==2)):
                    replicate=1

                dfTemp=Translation_Normal_tiled_PCA[(Translation_Normal_tiled_PCA["Cell"]==cell) 
                                                  & (Translation_Normal_tiled_PCA["Media"]==media) 
                                                  & (Translation_Normal_tiled_PCA["Time"]==time)
                                                  & (Translation_Normal_tiled_PCA["Replicate"]==replicate)]
                dfTemp.dropna(axis=0,inplace=True)
                dfTemp.reset_index(inplace=True)
                dfTemp.drop(['index'],axis=1,inplace=True)
                maxObjects=dfTemp.shape[0]
                maxCorrelation=20

                cellCorr=np.zeros((maxObjects,maxCorrelation))
                xPos=np.zeros(maxObjects)
                yPos=np.zeros(maxObjects)

                for i in range(0,maxObjects):
                    dfCurrentCell=dfTemp.iloc[[i]]
                    dfCellCorrCalc=dfCurrentCell
                    distanceToCells=np.sqrt((dfCurrentCell['AreaShape_Center_X'].values-dfTemp['AreaShape_Center_X'].values)**2+
                                                    (dfCurrentCell['AreaShape_Center_Y'].values-dfTemp['AreaShape_Center_Y'].values)**2)
                    tempList=distanceToCells[1:-1]
                    nearestCells=sorted(np.argpartition(tempList, maxCorrelation)[:maxCorrelation])
                    xPos[i]=dfCurrentCell['AreaShape_Center_X']
                    yPos[i]=dfCurrentCell['AreaShape_Center_Y']
                    for j in range(0,maxCorrelation):
                        dfCellCorrCalc=pd.concat([dfCellCorrCalc,dfTemp.iloc[[nearestCells[j]]]],ignore_index=True)
                        dfCellCorrCalc.drop(['Cell','Media','Time','ObjectNumber',
                                        'AreaShape_Center_X','AreaShape_Center_Y'],axis=1,inplace=True)
                        dfCellCorrCalcTranspose=dfCellCorrCalc.T
                        dfCellCorrMatrix=dfCellCorrCalcTranspose.corr()
                        if j > 0:
                            medianCorr=np.round(dfCellCorrMatrix[1:-1].median(axis=0),2)
                            cellCorr[i][j]=medianCorr[0]
                        else:
                            cellCorr[i][j]=np.round(dfCellCorrMatrix[0][1],2)

                # Add all median spatial correlations to final dataframe
                medianCorr=np.round(np.median(cellCorr,axis=1),2)
                dfTempCorr=pd.DataFrame(columns=['Cell type','Media type','Time point','Correlation'])
                dfTempCorr['Correlation']=medianCorr
                dfTempCorr['Cell type']=cell
                dfTempCorr['Media type']=media
                dfTempCorr['Time point']=time
                dfTempCorr['xPos']=xPos
                dfTempCorr['yPos']=yPos
                dfTempCorr['Replicate']=replicate

                Translation_Normal_SpatialCellCorr=pd.concat([Translation_Normal_SpatialCellCorr,dfTempCorr],ignore_index=True)   
                
                
#-------------------------------------------------------------------------------
# Tile images with PCA data for PPHN PAEC translation
#-------------------------------------------------------------------------------

# Setup conditions
cells=['PPHN']
times=[0,15,30,60,120,240,1440]
medias=['Vehicle','IGF1']
replicates=[1,2]

Translation_PPHN_SpatialCellCorr=pd.DataFrame(columns=['Cell type','Media type','Time point','Correlation', 'xPos','yPos'])

for time in times:
    for cell in cells:
        for media in medias:
            for replicate in replicates:
                
                # Fix missing replicates for PPHN data.
                # This won't be used in downstream analysis, it's just easier to do this way.
                if ((media=='Vehicle') & (time==120) & (replicate==2)):
                    replicate=1
                    
                if ((media=='IGF1') & (time==240) & (replicate==2)):
                    replicate=1

                dfTemp=Translation_PPHN_tiled_PCA[(Translation_PPHN_tiled_PCA["Cell"]==cell) 
                                                  & (Translation_PPHN_tiled_PCA["Media"]==media) 
                                                  & (Translation_PPHN_tiled_PCA["Time"]==time)
                                                  & (Translation_PPHN_tiled_PCA["Replicate"]==replicate)]
                dfTemp.dropna(axis=0,inplace=True)
                dfTemp.reset_index(inplace=True)
                dfTemp.drop(['index'],axis=1,inplace=True)
                maxObjects=dfTemp.shape[0]
                maxCorrelation=20

                cellCorr=np.zeros((maxObjects,maxCorrelation))
                xPos=np.zeros(maxObjects)
                yPos=np.zeros(maxObjects)

                for i in range(0,maxObjects):
                    dfCurrentCell=dfTemp.iloc[[i]]
                    dfCellCorrCalc=dfCurrentCell
                    distanceToCells=np.sqrt((dfCurrentCell['AreaShape_Center_X'].values-dfTemp['AreaShape_Center_X'].values)**2+
                                                    (dfCurrentCell['AreaShape_Center_Y'].values-dfTemp['AreaShape_Center_Y'].values)**2)
                    tempList=distanceToCells[1:-1]
                    nearestCells=sorted(np.argpartition(tempList, maxCorrelation)[:maxCorrelation])
                    xPos[i]=dfCurrentCell['AreaShape_Center_X']
                    yPos[i]=dfCurrentCell['AreaShape_Center_Y']
                    for j in range(0,maxCorrelation):
                        dfCellCorrCalc=pd.concat([dfCellCorrCalc,dfTemp.iloc[[nearestCells[j]]]],ignore_index=True)
                        dfCellCorrCalc.drop(['Cell','Media','Time','ObjectNumber',
                                        'AreaShape_Center_X','AreaShape_Center_Y'],axis=1,inplace=True)
                        dfCellCorrCalcTranspose=dfCellCorrCalc.T
                        dfCellCorrMatrix=dfCellCorrCalcTranspose.corr()
                        if j > 0:
                            medianCorr=np.round(dfCellCorrMatrix[1:-1].median(axis=0),2)
                            cellCorr[i][j]=medianCorr[0]
                        else:
                            cellCorr[i][j]=np.round(dfCellCorrMatrix[0][1],2)

                # Add all median spatial correlations to final dataframe
                medianCorr=np.round(np.median(cellCorr,axis=1),2)
                dfTempCorr=pd.DataFrame(columns=['Cell type','Media type','Time point','Correlation'])
                dfTempCorr['Correlation']=medianCorr
                dfTempCorr['Cell type']=cell
                dfTempCorr['Media type']=media
                dfTempCorr['Time point']=time
                dfTempCorr['xPos']=xPos
                dfTempCorr['yPos']=yPos
                dfTempCorr['Replicate']=replicate

                Translation_PPHN_SpatialCellCorr=pd.concat([Translation_PPHN_SpatialCellCorr,dfTempCorr],ignore_index=True)   

# %% [code]
%matplotlib inline

# Plot spatial maps of correlated cells for selected time points
# This generates Figures 8 and 9


sns.set_context("paper", font_scale=2)

times=[0,15,30,60,120,240,1440]
medias=['Vehicle','IGF1']
replicates=[1,2]

for media in medias:
    for time in times:
        for replicate in replicates:

            dfNull_plot=IF_Normal_NullCellCorr[((IF_Normal_NullCellCorr["Media type"]==media) 
                                        & (IF_Normal_NullCellCorr["Time point"]==time) 
                                        & (IF_Normal_NullCellCorr["Replicate"]==replicate))]

            dfNull_plot.dropna(axis=0,inplace=True)
            dfNull_plot.reset_index(inplace=True)
            dfNull_plot.drop(['index'],axis=1,inplace=True)
            null95percentile=np.round(np.quantile(dfNull_plot['Correlation'],q=0.95),2)

            dfSpatial_plot=IF_Normal_SpatialCellCorr[((IF_Normal_SpatialCellCorr["Media type"]==media) 
                                                      & (IF_Normal_SpatialCellCorr["Time point"]==time)
                                                      & (IF_Normal_SpatialCellCorr["Replicate"]==replicate))]

            dfSpatial_plot.dropna(axis=0,inplace=True)
            dfSpatial_plot.reset_index(inplace=True)
            dfSpatial_plot.drop(['index'],axis=1,inplace=True)
            #spatial95percentile=np.round(np.quantile(dfSpatial_plot['Correlation'],q=0.95),2)
            dfSpatial_plot['corrNullThreshold']=dfSpatial_plot['Correlation']>null95percentile
            #dfSpatial_plot['corrSpatialThreshold']=dfSpatial_plot['Correlation']>spatial95percentile
            dfSpatial_plot['corrNullThreshold']=np.multiply(dfSpatial_plot['corrNullThreshold'],1)
            #dfSpatial_plot['corrSpatialThreshold']=np.multiply(dfSpatial_plot['corrSpatialThreshold'],1)
            dfSpatial_plot['corrThreshold']=dfSpatial_plot['corrNullThreshold']#+dfSpatial_plot['corrSpatialThreshold']

            #dfSpatial_plot_dropped=dfSpatial_plot[dfSpatial_plot.corrThreshold > 0]

            g = sns.scatterplot(x="xPos", y="yPos", hue='corrThreshold',  data=dfSpatial_plot, legend=False,
                                s=(50),alpha=(0.4)).set_title('Normal PAEC; Media: '+str(media)+'; Time: '+str(time)+'; Replicate:'+str(replicate))
            plt.savefig('figure_07/Normal__Media_'+str(media)+'_Time_'+str(time)+'_Replicate+_'+str(replicate)+'.png', dpi=400)
            
for media in medias:
    for time in times:
        for replicate in replicates:

            dfNull_plot=IF_PPHN_NullCellCorr[((IF_PPHN_NullCellCorr["Media type"]==media) 
                                        & (IF_PPHN_NullCellCorr["Time point"]==time) 
                                        & (IF_PPHN_NullCellCorr["Replicate"]==replicate))]

            dfNull_plot.dropna(axis=0,inplace=True)
            dfNull_plot.reset_index(inplace=True)
            dfNull_plot.drop(['index'],axis=1,inplace=True)
            null95percentile=np.round(np.quantile(dfNull_plot['Correlation'],q=0.95),2)

            dfSpatial_plot=IF_PPHN_NullCellCorr[((IF_PPHN_NullCellCorr["Media type"]==media) 
                                                      & (IF_PPHN_NullCellCorr["Time point"]==time)
                                                      & (IF_PPHN_NullCellCorr["Replicate"]==replicate))]

            dfSpatial_plot.dropna(axis=0,inplace=True)
            dfSpatial_plot.reset_index(inplace=True)
            dfSpatial_plot.drop(['index'],axis=1,inplace=True)
            #spatial95percentile=np.round(np.quantile(dfSpatial_plot['Correlation'],q=0.95),2)
            dfSpatial_plot['corrNullThreshold']=dfSpatial_plot['Correlation']>null95percentile
            #dfSpatial_plot['corrSpatialThreshold']=dfSpatial_plot['Correlation']>spatial95percentile
            dfSpatial_plot['corrNullThreshold']=np.multiply(dfSpatial_plot['corrNullThreshold'],1)
            #dfSpatial_plot['corrSpatialThreshold']=np.multiply(dfSpatial_plot['corrSpatialThreshold'],1)
            dfSpatial_plot['corrThreshold']=dfSpatial_plot['corrNullThreshold']#+dfSpatial_plot['corrSpatialThreshold']

            #dfSpatial_plot_dropped=dfSpatial_plot[dfSpatial_plot.corrThreshold > 0]
            
            g = sns.scatterplot(x="xPos", y="yPos", hue='corrThreshold',  data=dfSpatial_plot, legend=False,
                                s=(50),alpha=(0.4)).set_title('PPHN PAEC; Media: '+str(media)+'; Time: '+str(time)+'; Replicate:'+str(replicate))
            plt.savefig('figure_07/PPHN_Media_'+str(media)+'_Time_'+str(time)+'_Replicate+_'+str(replicate)+'.png', dpi=400)

# %% [code]
# Plot median percentage of cells that are spatially correlated
# This generates Figure 9

times=[0,15,30,60,120,240,1440]
medias=['Vehicle','IGF1']
replicates=[1,2]

dfPerct=pd.DataFrame(columns=['Media','Time','Replicate','corrCells'])

for media in medias:
    for time in times:
        for replicate in replicates:

            dfNull_perct=dfNullCellCorr[((dfNullCellCorr["Media type"]==media) 
                           & (dfNullCellCorr["Time point"]==time) 
                           & (dfNullCellCorr["Replicate"]==replicate))]

            dfNull_perct.dropna(axis=0,inplace=True)
            dfNull_perct.reset_index(inplace=True)
            dfNull_perct.drop(['index'],axis=1,inplace=True)
            null95percentile=np.round(np.quantile(dfNull_perct['Correlation'],q=0.95),2)

            dfSpatial_perct=dfSpatialCellCorr[((dfSpatialCellCorr["Media type"]==media) 
                                              & (dfSpatialCellCorr["Time point"]==time)
                                              & (dfSpatialCellCorr["Replicate"]==replicate))]

            dfSpatial_perct.dropna(axis=0,inplace=True)
            dfSpatial_perct.reset_index(inplace=True)
            dfSpatial_perct.drop(['index'],axis=1,inplace=True)
            dfSpatial_perct['corrNullThreshold']=dfSpatial_perct['Correlation']>null95percentile
            
            corrPerct=(dfSpatial_perct['corrNullThreshold'].sum()/dfSpatial_perct.shape[0])
            d = {'Media': [media],'Time': [time], 'Replicate': [replicate], 'corrCells': [corrPerct]}
            dfTempCellCorr=pd.DataFrame(data=d)
            dfPerct=pd.concat([dfPerct,dfTempCellCorr],ignore_index=True)

# %% [code]
sns.set_context("paper", rc={"font.size":8,"axes.titlesize":8,"axes.labelsize":5})   

g = sns.catplot(x = "Time", y="corrCells", hue="Media", ci=None, data=dfPerct, kind="point", scale=2)
axes = g.axes
axes[0,0].set_title("Normal PAEC",fontsize=20)
axes[0,0].set_ylim(-.05,.25)
axes[0,0].set_xlabel("Time",fontsize=20)
axes[0,0].set_ylabel("Percent PAEC with \nsignificant spatial correlation",fontsize=20)
axes[0,0].tick_params(labelsize=15)
plt.legend(fontsize='x-large', title_fontsize='40')
```
