## Supplemental code for "High throughput imaging identifies a spatially localized response of primary fetal pulmonary artery endothelial cells to insulin-like growth factor 1 treatment": generate_supplemental_figures.ipynb

```
# %% [code]
# import libraries for analysis and plotting
import pandas as pd
import seaborn as sns
import numpy as np
import scipy as sp
import random
import matplotlib.pyplot as plt
from sklearn.preprocessing import StandardScaler
from sklearn.decomposition import PCA
from statsmodels import robust
import warnings
warnings.filterwarnings('ignore')

# This library must be downloaded and installed into the active environment 
# from: https://github.com/webermarcolivier/statannot
# from statannot import add_stat_annotation 

# %% [code]
#Import dose response data
df_dose=pd.read_csv("ensemble/dose.csv")

# %% [code]
#Plot and save dose response data
ax_dose = sns.catplot("Cell", "Cells", "Media (ng/ml)", kind='strip', data=df_dose, dodge=True, s=15, aspect=2)
axes_dose=ax_dose.axes

axes_dose[0,0].set_yticks([80000, 120000, 160000])

sns.despine(offset=5, trim=False)
fig = ax_dose.fig
fig.savefig('dose.png', dpi=600) 

# %% [code]
#Import Axitinib data
df_Ax=pd.read_csv("ensemble/axitinib.csv")

#Create individual data structures for statistical tests
df_Ax_Vehicle=df_Ax[df_Ax['Media'] == 'Vehicle']
df_Ax_IGF1=df_Ax[df_Ax['Media'] == 'IGF1']
df_Ax_VEGF=df_Ax[df_Ax['Media'] == 'VEGF']

# %% [code]
#Plot and save axitinib data
ax_axitinib = sns.catplot("Cell", "Cells", "Inhibitor", col="Media", data=df_Ax, kind="strip", dodge=True, s=15, sharey=False, aspect=0.9)
sns.set_context("paper", font_scale=3)
axes_axitinib=ax_axitinib.axes

add_stat_annotation(axes_axitinib[0,0], data=df_Ax_Vehicle, x="Cell", y="Cells", hue="Inhibitor",
                    boxPairList=[(("Normal", "None"), ("Normal", "Axitinib")),
                                (("PPHN", "None"), ("PPHN", "Axitinib"))],
                    test='Kruskal', textFormat='star', loc='inside', verbose=2)

add_stat_annotation(axes_axitinib[0,1], data=df_Ax_IGF1, x="Cell", y="Cells", hue="Inhibitor",
                    boxPairList=[(("Normal", "None"), ("Normal", "Axitinib")),
                                (("PPHN", "None"), ("PPHN", "Axitinib"))],
                    test='Kruskal', textFormat='star', loc='inside', verbose=2)

add_stat_annotation(axes_axitinib[0,2], data=df_Ax_VEGF, x="Cell", y="Cells", hue="Inhibitor",
                    boxPairList=[(("Normal", "None"), ("Normal", "Axitinib")),
                                (("PPHN", "None"), ("PPHN", "Axitinib"))],
                    test='Kruskal', textFormat='star', loc='inside', verbose=2)

axes_axitinib[0,0].set_yticks([100000, 200000])
axes_axitinib[0,1].set_yticks([100000, 200000, 300000])
axes_axitinib[0,2].set_yticks([100000, 200000, 300000])

sns.despine(offset=5, trim=False)
fig = ax_axitinib.fig
fig.savefig('axitinib.png', dpi=600) 

# %% [code]
#Import PPP treatment
df_PPP=pd.read_csv("ensemble/PPP.csv")

#Create individual data structures for statistical tests
df_PPP_Vehicle=df_PPP[df_PPP['Media'] == 'Vehicle']
df_PPP_IGF1=df_PPP[df_PPP['Media'] == 'IGF1']
df_PPP_VEGF=df_PPP[df_PPP['Media'] == 'VEGF']

# %% [code]
#Plot and save PPP data
ax_PPP = sns.catplot("Cell", "Cells", "Inhibitor", col="Media", data=df_PPP, kind="strip", dodge=True, s=15, sharey=False)
sns.set_context("paper", font_scale=3)
axes_PPP=ax_PPP.axes


add_stat_annotation(axes_PPP[0,0], data=df_PPP_Vehicle, x="Cell", y="Cells", hue="Inhibitor",
                    boxPairList=[(("Normal", "None"), ("Normal", "PPP"))],
                    test='Kruskal', textFormat='star', loc='inside', verbose=2)

add_stat_annotation(axes_PPP[0,1], data=df_PPP_IGF1, x="Cell", y="Cells", hue="Inhibitor",
                    boxPairList=[(("Normal", "None"), ("Normal", "PPP"))],
                    test='Kruskal', textFormat='star', loc='inside', verbose=2)

add_stat_annotation(axes_PPP[0,2], data=df_PPP_VEGF, x="Cell", y="Cells", hue="Inhibitor",
                    boxPairList=[(("Normal", "None"), ("Normal", "PPP"))],
                    test='Kruskal', textFormat='star', loc='inside', verbose=2)

axes_PPP[0,0].set_yticks([100000, 150000])
axes_PPP[0,1].set_yticks([100000, 150000, 200000, 250000])
axes_PPP[0,2].set_yticks([100000, 150000, 200000, 250000])

sns.despine(offset=5, trim=False)
fig = ax_PPP.fig
fig.savefig('PPP.png', dpi=600) 

# %% [code]
#import Rapamyocin treatment
df_RP=pd.read_csv("ensemble/RP.csv")

#Create individual data structures for statistical tests
df_RP_Vehicle=df_RP[df_RP['Media'] == 'Vehicle']
df_RP_IGF1=df_RP[df_RP['Media'] == 'IGF1']
df_RP_VEGF=df_RP[df_RP['Media'] == 'VEGF']

# %% [code]
#Plot and save RP data
ax_RP = sns.catplot("Cell", "Cells", "Inhibitor", col="Media", data=df_RP, kind="strip", dodge=True, s=15, sharey=True)
sns.set_context("paper", font_scale=3)
axes_RP=ax_RP.axes

add_stat_annotation(axes_RP[0,0], data=df_RP_Vehicle, x="Cell", y="Cells", hue="Inhibitor",
                    boxPairList=[(("Normal", "None"), ("Normal", "Rapamycin (low)")),
                                (("Normal", "None"), ("Normal", "Rapamycin (high)"))],
                    test='Kruskal', textFormat='star', loc='inside', verbose=2)

add_stat_annotation(axes_RP[0,1], data=df_RP_IGF1, x="Cell", y="Cells", hue="Inhibitor",
                    boxPairList=[(("Normal", "None"), ("Normal", "Rapamycin (low)")),
                                (("Normal", "None"), ("Normal", "Rapamycin (high)"))],
                    test='Kruskal', textFormat='star', loc='inside', verbose=2)

add_stat_annotation(axes_RP[0,2], data=df_RP_VEGF, x="Cell", y="Cells", hue="Inhibitor",
                    boxPairList=[(("Normal", "None"), ("Normal", "Rapamycin (low)")),
                                (("Normal", "None"), ("Normal", "Rapamycin (high)"))],
                    test='Kruskal', textFormat='star', loc='inside', verbose=2)

sns.despine(offset=5, trim=False)
fig = ax_RP.fig
fig.savefig('RP.png', dpi=600) 
```
