## Supplemental code for "High throughput imaging identifies a spatially localized response of primary fetal pulmonary artery endothelial cells to insulin-like growth factor 1 treatment": preprocess_raw_data.ijm

```
//Script to convert sCMOS data to photons before running NCS correction in MATLAB
//Conversion values dervied from camera datasheets (C1,C2 & C3,C4 obtained on different cameras)
//Shepherd laboratory
//2019.06.14

path = getDirectory("Choose a Directory"); 
filename = getFileList(path); 
photonCorrectPath = path + "photonCorrectPath" + File.separator; 
if (File.exists(photonCorrectPath)) 
	exit("Destination directory already exists; remove it and then run this macro again"); 
File.makeDirectory(photonCorrectPath); 

setBatchMode(true)
for (i=0; i<filename.length; i++) { 
	if(endsWith(filename[i], ".tif")) { 
		open(path+filename[i]); 
		run("Properties...", "channels=4 slices=51 frames=1 unit=nm pixel_width=162.5 pixel_height=162.5 voxel_depth=350.0");
		setMinAndMax(0, 65535);
		run("Split Channels");
		list = getList("image.titles");
		for (j=0; j<=3;j++) {
			selectWindow(list[j]);
			if(startsWith(list[j],"C1")) {
				run("Multiply...", "value=1 stack");
				run("Subtract...", "value=100 stack");
			}
			if(startsWith(list[j],"C2")) {
				run("Multiply...", "value=0.78 stack");
				run("Subtract...", "value=78 stack");
			}
			if(startsWith(list[j],"C3")) {
				run("Multiply...", "value=0.730 stack");
				run("Subtract...", "value=73 stack");
			}
			if(startsWith(list[j],"C4")) {
				run("Multiply...", "value=0.820 stack");
				run("Subtract...", "value=82 stack");
			}
		}
		run("Merge Channels...","c1=C1-"+filename[i]+" c2=C2-"+filename[i]+" c3=C3-"+filename[i] +" c4=C4-"+filename[i]+" create");
		saveAs("tif", photonCorrectPath+filename[i]);
		close();
		ok=File.delete(path+filename[i]);
	} 
}
setBatchMode(false)
;
```
