## Supplemental code for "High throughput imaging identifies a spatially localized response of primary fetal pulmonary artery endothelial cells to insulin-like growth factor 1 treatment": process_deconvolved_data.ijm

```
//Script to flat field correct and make z projection after NCS denoising and batch deconvolution
//Shepherd laboratory
//2019.06.14

path = getDirectory("Choose root directory"); 
deconpath= path + "decon" + File.separator; 
filename = getFileList(deconpath); 
newDir = path + "Zprojection" + File.separator; 
if (File.exists(newDir)) 
	exit("Destination directory already exists; remove it and then run this macro again"); 
File.makeDirectory(newDir); 
for (i=0; i<filename.length; i++) { 
	if(endsWith(filename[i], ".tif")) { 
		open(deconpath+filename[i]); 
		imageTitle=getTitle();
		run("BaSiC ", "processing_stack=["+imageTitle+"] flat-field=None dark-field=None shading_estimation=[Estimate shading profiles] shading_model=[Estimate flat-field only (ignore dark-field)] setting_regularisationparametes=Automatic temporal_drift=[Replace with zero] correction_options=[Compute shading and correct images] lambda_flat=0.50 lambda_dark=0.50");
		selectWindow("Corrected:"+imageTitle);
		run("Z Project...", "projection=[Max Intensity]");
		selectWindow("MAX_Corrected:"+imageTitle);
		saveAs("tif", newDir + imageTitle);
		while (nImages>0) { 
          selectImage(nImages); 
          close(); 
      	} 
      	ok=File.delete(deconpath+filename[i]);
	} 
}
```
